# A TRANSPARENT WHEEL-BASED PLATFORM FOR LOCOMOTION-ON-DEMAND AND MULTI-VIEW BODY AND FACIAL KINEMATICS IN HEAD-FIXED MICE

**DOI:** 10.64898/2025.12.26.696644

**Authors:** Pratik S Paranjape, Tahoura Mohammadi Ghohaki, Samsoon Inayat

## Abstract

Understanding how the brain transforms sensory input and internal state into coordinated action requires behavioral paradigms that provide precise, multimodal measurements of movement and arousal while remaining compatible with neural recording techniques. Here we present a modular behavioral platform that enables stimulus-evoked locomotion in head-fixed mice using a transparent running wheel combined with air-stream stimulation. The design provides direct ventral access for imaging paw movements while simultaneously capturing body kinematics, facial motion, and eye-related signals from multiple camera views. The system integrates Arduino-based stimulus control, rotary encoder measurements, Raspberry Pi–based videography, and LED-based visual markers for temporal alignment across independently acquired data streams. Using a proof-of-principle dataset from well-trained animals, we show that brief air delivery reliably induces structured locomotion with reproducible trial timing. Optical-flow–based motion metrics and DeepLabCut pose estimation reveal robust, stimulus-locked increases in paw, limb, and facial movements during air-on epochs relative to air-off periods. LED-based event markers enable consistent identification of air-on and air-off epochs across video streams despite differences in sampling rates. Together, these features provide a flexible framework for studying stimulus-driven locomotion and multi-view behavioral dynamics under head fixation, with straightforward compatibility for integration with neural imaging and electrophysiology recording approaches.

**Significance Statement:** Quantifying how the brain generates coordinated movement requires behavioral paradigms that provide precise, multi-view measurements while remaining compatible with neural recording techniques. We introduce a low-cost, modular behavioral platform that enables locomotion-on-demand in head-fixed mice using a transparent running wheel, allowing simultaneous visualization of ventral paw movements, facial dynamics, and eye/pupil-related signals. By combining stimulus-induced locomotion with synchronized multi-camera videography and open-source analysis pipelines, this system overcomes key limitations of existing head-fixed locomotion assays that rely on prolonged reward-based training or provide only limited behavioral readouts.

## Introduction

Coordination and motor control are primary functions of the brain that enable animals to navigate their environment and survive. The motor-control problem solved by the brain is highly complex: movements involve multiple degrees of freedom, and the same action can be achieved through different combinations of muscles ^1–3^. This redundancy means that understanding how the brain generates coordinated movement requires precise measurement of behavior at multiple levels, from whole-body and limb kinematics to facial and autonomic indicators of arousal ^4–6^. To link these components with neural activity, it is essential to develop advanced experimental setups capable of capturing motor output and enabling measurement of internal state (e.g., via pupillometry) with high temporal and spatial precision ^7–9^. Developing such comprehensive systems is a critical step toward enabling quantitative behavioral characterization, which is the goal of the present study.

Behavioral paradigms have evolved over the past century from early ethological observations to laboratory research employing conditioning boxes, mazes, and sophisticated head-fixed tasks incorporating virtual reality, videography, electrophysiology, and brain imaging ^10–16^. Although head fixation involves unnatural restraint, it enables spatially precise imaging of brain structure and activity ^17–19^. Most experimental rigs, however, quantify only limited aspects of behavior, for example belt speed as a proxy for running behavior on a conveyor belt or kinematics of a restricted set of body parts (limbs, whiskers, or eyes), typically using a single video stream, thereby constraining the range of investigative questions that can be addressed ^19–22^. Furthermore, many behavioral paradigms rely on extensive reward-based conditioning, in which water-deprived animals receive water as a reward, and training times often span several weeks, limiting throughput and experimental efficiency ^23,24^. Stimulus-induced locomotion paradigms, such as air-stream–based approaches, offer an alternative by evoking locomotion on demand without prolonged conditioning and have been demonstrated primarily using treadmill or linear conveyor-belt systems ^18,19^. However, these approaches have not been systematically implemented or validated using transparent running wheels that would permit unobstructed, multi-angle videography of the animal during locomotion. An important gap therefore remains in determining whether stimulus-induced locomotion can be combined with a wheel-based, visually accessible platform to enable simultaneous, high–temporal resolution measurement of whole-body kinematics, facial dynamics, and pupillometry in head-fixed mice. Bridging this gap requires integration of behavioral motivation, precise stimulus control, and synchronized multi-view recording within a single experimental framework. This study presents a novel behavioral setup that enables locomotion-on-demand with simultaneous measurement of body and facial kinematics together with ocular measurements in head-fixed mice. The system is low-cost and modular, constructed primarily from readily available components, including Arduino and Raspberry Pi boards, and is accompanied by detailed design schematics, assembly guide, list of materials, and open-source code for stimulus control, data acquisition, and analyses. With this system, we present the successful implementation of the air-induced running paradigm (AIR), which enables stable head-fixed locomotion in response to the application of air to the animal’s back while simultaneously recording body, facial, and ocular dynamics. We further demonstrate behavioral analysis using optical flow-based motion analysis and DeepLabCut-based pose estimation.

## Ethics Statement, Experimental model, and subject details

This study is reported in accordance with the ARRIVE guidelines (https://arriveguidelines.org). All animal procedures were performed in compliance with protocols approved by the Institutional Animal Care and Use Committee (IACUC) of the University of Nevada, Las Vegas, and were conducted in accordance with the guidelines for the ethical use of animals established by the National Institutes of Health and relevant institutional regulations. An adult male C57BL/6J-Tg (Thy1-GCaMP6s) GP4.3Dkim/J mouse (∼14 months old; body weight 32.3 g at the time of experiments) was used in this study as a representative animal. Head-fixation surgery was performed at ∼13 months of age. In addition, three adult male C57BL/6J mice (approximately 8 months old at the time of behavioral testing) were included to assess cross-animal reproducibility of the AIR-wheel paradigm. Body weight at the time of experimentation for these animals ranged from 29 – 33 g. Head-fixation surgeries for these animals were performed ∼2 months prior to behavioral testing. All animals were obtained from the Jackson Laboratory and housed under standard laboratory conditions at the University of Nevada, Las Vegas animal facility. At the conclusion of the study, the mice were euthanized using carbon dioxide (CO₂) inhalation followed by cervical dislocation, in accordance with protocols approved by the IACUC, and conducted in compliance with the guidelines for the ethical use of animals established by the National Institutes of Health and relevant institutional regulations. CO₂ was delivered from a compressed gas cylinder at a controlled displacement rate of approximately 3.4 L/min, with room air present at the start of exposure. Gas flow was maintained until loss of consciousness and continued for at least 1-2 minutes following cessation of respiration. Cervical dislocation was subsequently performed by trained personnel as a secondary physical method to ensure death.

## Methods

### Housing and husbandry

Animals were housed in a temperature- and humidity-controlled facility under a standard 12 h light/12 h dark cycle with ad libitum access to food and water. Mice were housed in standard laboratory cages in accordance with institutional guidelines. Animals were monitored daily following surgery and during behavioral training for general health, body condition, and signs of distress. No unexpected adverse events occurred. Humane endpoints were defined in accordance with institutional guidelines and included signs of persistent distress, significant weight loss, or impaired mobility; none were reached.

### Surgery

The procedure was carried out using sterile technique. Animals received a subcutaneous injection of 0.2 mL of a sterile 0.9% saline solution before surgery. Anesthesia was induced and maintained with isoflurane (2–3% for induction; 1–1.5% for maintenance), and the animal was placed on a heating pad. Following induction of anesthesia, the mouse was head-fixed in a stereotaxic apparatus (Harvard Apparatus, Inc). A midline incision was made to expose the skull. The skull surface was cleaned and dried, and a custom-designed headplate was positioned and secured to the skull using dental cement (C&B METABOND®). The headplate was fabricated from black polylactic acid (PLA) material (Official Creality Hyper PLA Filament 1.75mm) using a 3D printer (Creality K1C 3D) and designed to provide stable and repeatable head fixation during behavioral experiments.

After implantation, the skin surrounding the headplate was sealed using dental acrylic as needed. Postoperative care included subcutaneous administration of meloxicam (5 mg/kg) for analgesia for 2–3 days following surgery. Animals were allowed to recover fully before habituation to head fixation and subsequent behavioral testing. Behavioral experiments were conducted when the animals were ∼14 months old.

### Behavioral Setup

The details of the mechanical and electronic hardware (List of Materials), along with associated design, and assembly guide are provided in the Supplementary Materials.

#### Mechanical Hardware

The behavioral apparatus was built on a rigid aluminum base plate (approximately 8 × 12 inches, 0.5-inch thickness), which served as the primary mechanical support for all components. A central hole was drilled into the base plate to mount a vertical cylindrical aluminum rod, which functioned as the main structural axis of the setup. All major mechanical elements, including the running wheel, head-fixation assembly, camera mounts, and air-delivery components, were attached either to this rod or the base plate using standard laboratory clamps and rods (Fig. 1).

**Figure 1.**
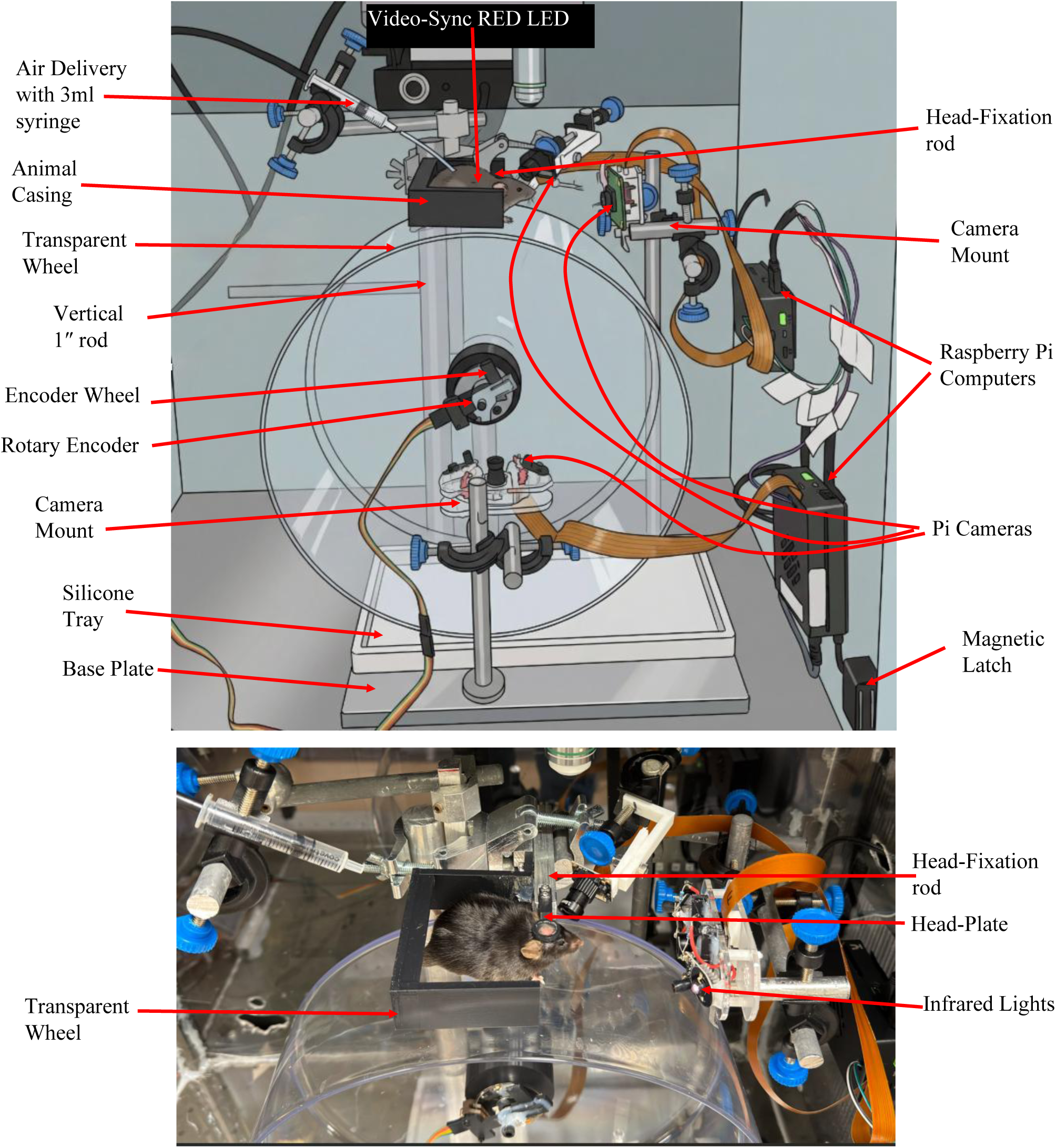
Overview of the Behavioral Apparatus. Schematic (top) and a photograph (bottom) of the behavioral setup illustrating the major mechanical and electronic components. The system is built around a rigid aluminum base plate supporting a vertical cylindrical rod that serves as the primary structural axis. A transparent running wheel is mounted on the rod and instrumented with a rotary encoder for measurement of wheel position and speed. A removable silicone tray positioned beneath the wheel collects feces and urine during experiments. The animal is head-fixed using a custom headplate clamp attached to the vertical rod. Air delivery is achieved via a syringe-based nozzle positioned above the animal’s back. Multiple cameras are mounted around the apparatus for bottom-view, facial, and pupil imaging and are connected to Raspberry Pi computers for synchronized video acquisition.

A non-motorized transparent running wheel (Bucatstate Hamster Wheel) was mounted on the vertical rod using the manufacturer provided attachment. For the wheel, the shaft rod itself was fixed, while the wheel rotated freely around it. This configuration allowed unobstructed visual access to the animal from below and the sides while maintaining mechanical stability during locomotion. The entire apparatus was enclosed within a casing constructed from 1/4-inch acrylic sheets, forming a transparent enclosure around the animal to provide physical containment while preserving visibility for videography. A removable silicone tray was placed on the base plate directly beneath the transparent running wheel to collect feces and urine during experiments, facilitating hygiene and rapid cleaning between sessions.

Head fixation was achieved by attaching a custom headplate clamp assembly to the same vertical rod. A rectangular aluminum bar with a square cross-section was mounted using clamps and served as the support for the headplate holder. The headplate holder incorporated two threaded holes to secure the implanted headplate using screws. This arrangement allowed precise and repeatable positioning of the animal’s head relative to the running wheel and cameras. A 3D printed rigid animal casing was positioned around the head-fixed region of the wheel to restrict lateral movement and prevent the animal from reaching the edges of the running wheel during locomotion.

Multiple cameras were mounted around the apparatus using standard acrylic camera holders supplied with the cameras. These holders were secured to rods and clamps attached to the aluminum base plate, enabling flexible positioning and alignment of cameras for bottom-view, facial, and pupil imaging.

Wheel position and rotation were monitored using a rotary encoder mounted directly to the running wheel via a custom-designed, 3D-printed fixture. In this configuration, the encoder wheel rotated with the transparent running wheel, while the encoder body was fixed to the stationary horizontal shaft rod. This design enabled accurate measurement of wheel position and speed without interfering with wheel rotation or animal movement.

Finally, the vertical rod also supported the air-delivery tubing. Pressurized air was supplied from a laboratory wall outlet (∼70 PSI) and routed through an inline pressure regulator to reduce the operating pressure to ∼10 PSI at the setup. Regulated air was delivered to the animal via flexible tubing connected to a 3 mL syringe mounted on the mechanical frame, which served as a stable and precisely positionable air nozzle. Airflow to the syringe was gated by a solenoid valve placed in-line with the tubing and controlled electronically via a solid-state relay driven by the Arduino (see below). This configuration enabled rapid, reproducible onset and offset of the air stream with minimal mechanical vibration while allowing fine adjustment of both air pressure and nozzle positioning relative to the animal.

#### Electronic Hardware

The electronic hardware integrates behavioral control, kinematic measurement, and synchronized data acquisition across multiple devices. Locomotion was quantified using a rotary quadrature encoder (Broadcom / Avago 3 Channel 1000CPR) mounted to the running wheel, providing two digital quadrature outputs (Channel A and Channel B). These encoder signals were routed in parallel to an Arduino microcontroller (Mega 6250) for real-time behavioral control and to a National Instruments USB-6259 data acquisition (DAQ) card for high-resolution recording. On the DAQ, encoder signals were acquired using hardware counters and sampled in MATLAB at 5 kHz, allowing precise reconstruction of wheel position and running speed (Fig. 2).

**Figure 2.**
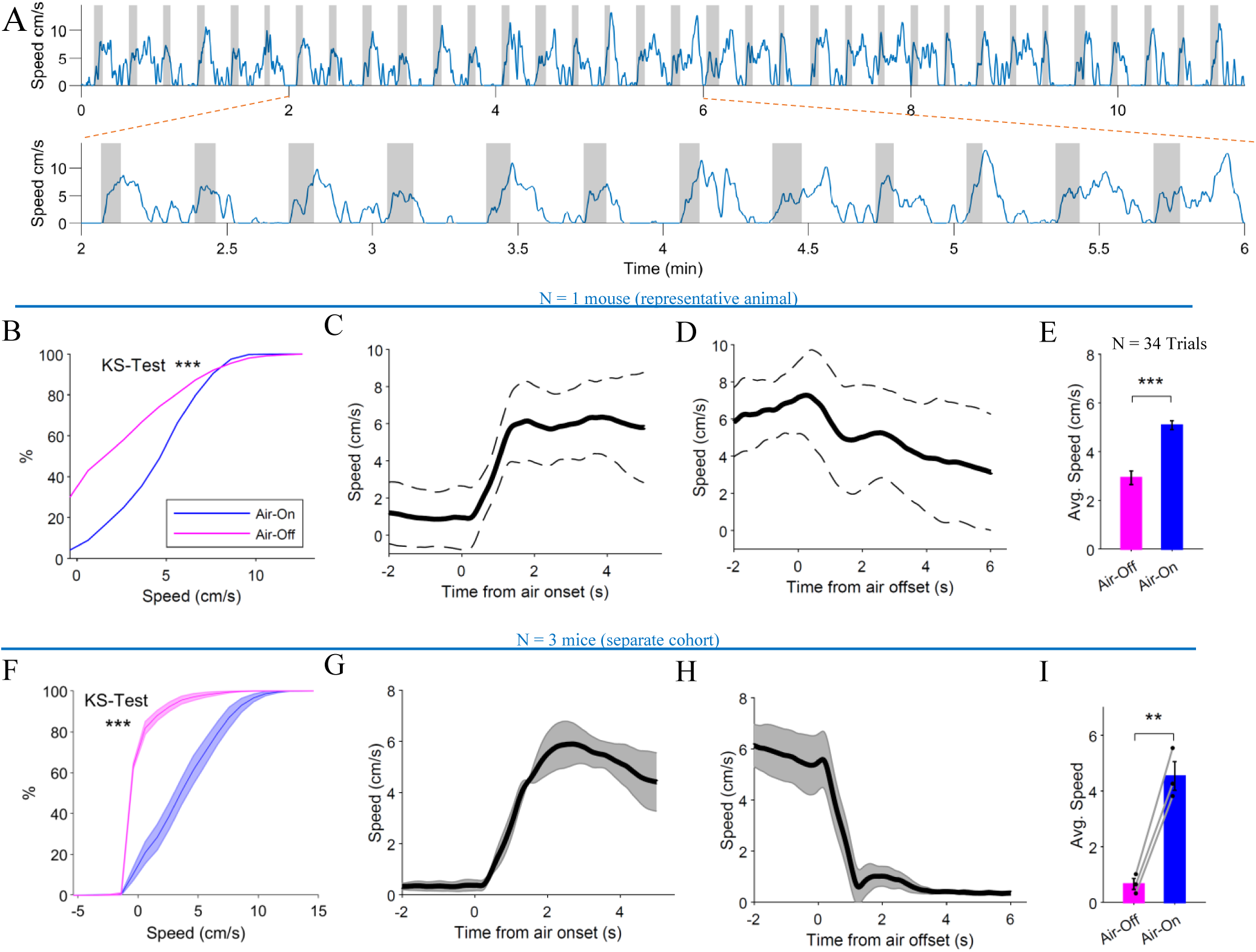
Air-induced locomotion produces reliable and trial-structured running behavior on a transparent wheel. **(A)** Continuous recording of wheel-derived running speed (blue) and air stimulus timing (orange) during an example ∼11-minute session comprising 34 trials. Air application reliably elicited increases in running speed, which decreased following air offset. **(B)** Cumulative distributions of running speed during air-on (blue) and air-off (magenta) periods across the session. The distributions were significantly different (Kolmogorov–Smirnov test, *** p < 0.001), indicating a separation between locomotor states. **(C)** Trial-aligned running speed relative to air onset (time 0 s). Solid line indicates the mean speed across trials; dashed lines indicate ±1 standard deviation. Running speed increased rapidly following air onset and reached a stable plateau during air application. **(D)** Trial-aligned running speed relative to air offset (time 0 s). Termination of the air stimulus was followed by a gradual reduction in running speed, demonstrating precise stimulus-locked control of locomotion. **(E)** Mean running speed computed on a per-trial basis during air-on and the immediately preceding air-off periods (N = 34 trials). Speeds were significantly higher during air-on epochs compared with air-off epochs (paired t-test and Wilcoxon signed-rank test, *** p < 0.001 for both). **(F – I)** Same as B-E respectively but for a separate cohort of N = 3 mice. Lines and bar graphs represent mean over trials and animals and shaded regions represent the standard error of the mean across animals. Individual dots in (I) indicate mean values for individual animals (** p < 0.01, paired t-test; Wilcoxon signed-rank test not significant).

The Arduino continuously monitored the encoder signals to implement distance-based control of the behavioral paradigm (see below). When predefined criteria were met, the Arduino generated a TTL output that drove a solid-state relay, which in turn actuated a solenoid valve controlling the air stream used to induce locomotion. The same air-control TTL signal was also routed to the USB-6259 and recorded at 5 kHz, enabling precise temporal alignment between locomotion, stimulus delivery, and behavioral state. Behavioral execution was gated by dual control inputs: a software-generated TTL signal from the USB-6259 (digital output) and a manual hardware switch. Both signals were required to be high for the behavioral paradigm to run, allowing flexible control and safe manual intervention during experiments. The DAQ-generated TTL control signal was additionally distributed to Raspberry Pi computers via GPIO pin 17. On each Raspberry Pi, custom Python acquisition scripts monitored this GPIO line and initiated video recording only when the trigger signal was high.

Two Raspberry Pi systems were used for videographic acquisition. One Raspberry Pi recorded a single camera providing a ventral (bottom) view of the animal and running wheel, while a second Raspberry Pi controlled two cameras simultaneously for facial and pupil imaging. All cameras, except the pupil camera, were illuminated using infrared (IR) light sources to enable video acquisition in complete darkness, as all training and testing were performed without visible illumination. Camera resolution (1440 x 1080 for bottom and face cameras, 320 x 240 for pupil camera), frame rate (60Hz), and acquisition parameters were configured via custom Python code running on the Raspberry Pi systems.

Video recording commenced automatically upon receipt of a TTL trigger signal. When MATLAB-based acquisition on the USB-6259 was terminated, the trigger signal was driven low, stopping both the behavioral paradigm and video acquisition in a synchronized manner. For this reason, the total duration of the DAQ recording was used to estimate the effective frame rates for the individual video streams (as compared to the nominal 60Hz frame rate set in the software) and were used for subsequent analysis for setting a time axis and alignment of the signals around air onset and offset events.

Video-to-behavior synchronization was further validated using an LED-based timing marker. A visible red LED mounted within the camera field of view was driven by the same TTL signal that controlled air delivery. As a result, LED illumination provided a direct visual indicator of air onset and offset within the video streams, enabling precise alignment of videographic, behavioral, and kinematic data during offline analysis.

### Behavioral Paradigm: Air-Induced Running on A Wheel

Animals were habituated to head fixation over 3-18 consecutive days depending on the cohort. During habituation, mice were head-fixed once or twice per day, with session durations gradually increased from 5 to 30 minutes to acclimate animals to restraint and the behavioral apparatus. Following habituation, mice were trained to run on the transparent running wheel while head fixed. All movement was generated voluntarily by the animal as the wheel is non-motorized. Training and testing were performed in the absence of visual cues, and the wheel surface was cleaned between sessions.

Animals underwent approximately up to two weeks of task training before experimental recordings. Training sessions lasted ∼20 minutes per day and were conducted at the same time each day. During this period, mice learned to walk on the wheel while head-fixed and to initiate locomotion in response to brief air stimulation directed toward the lower back. For the additional cohort of three wild-type mice used to assess cross-animal reproducibility, the number of training days prior to testing was 12, 2, and 4 days, respectively.

Air delivery was controlled by a solenoid valve driven by an Arduino-based system and supplied via the building’s compressed air line through a pressure regulator. The air stream served as a movement-inducing stimulus rather than an aversive reinforcer. On each trial, air onset initiated a running epoch during which the animal was required to run a predefined distance on the wheel, measured using the rotary encoder attached to the wheel. Once the target distance was reached, the air stream was automatically terminated, marking the end of the trial. Trials were separated by an intertrial interval of 15 s ^18,19^.

Training and testing both were carried out in a dark environment. Training followed a progressive distance schedule. The required running distance initially started at short distances ∼5cm and was gradually increased across trials. In the final configuration used for testing, animals were required to run a fixed distance of 25 cm to terminate the air stimulus. If the animal completed the target distance with sufficient and continuous locomotion, the trial was considered successful. Animals were trained until they reliably completed the required distance across the majority of trials within a session. Following training, the same paradigm was used during data collection sessions for behavioral and videographic analysis.

### Experimental Design

This study employed a within-subject experimental design in which head-fixed mice served as their own controls. Behavioral and kinematic measures obtained during air-on epochs were compared with those obtained during air-off epochs within the same recording sessions. The experimental unit for primary behavioral comparisons was the individual animal, with within-session analyses performed at the trial level. For the additional cohort (N = 3 mice), cross-animal consistency was assessed using repeated-measures ANOVA to evaluate reproducibility of air-evoked locomotion. Randomization was not used, as this study employed a within-subject design. Blinding was not performed during allocation, experimental conduct, outcome assessment, or data analysis due to the technical and proof-of-principle focus of the study.

### Inclusion and exclusion criteria

No a priori inclusion or exclusion criteria were defined for animals, trials, or data points. All recorded trials and data points were included in the analyses, and no exclusions were made.

### Data Analysis

Behavioral and control signals were analyzed using custom-written scripts in MATLAB (MathWorks). Encoder signals acquired via the USB-6259 were processed to extract wheel position, distance traveled, and running speed for each trial. Behavioral events, including air onset and offset, were aligned to encoder-derived kinematic measures using recorded TTL synchronization signals. In addition to hardware-based TTL synchronization, air onset and offset were independently identified directly from the video recordings using an LED timing marker positioned within the camera field of view. The LED was driven by the same control signal as the air-delivery solenoid, allowing air-related events to be extracted directly from the videos and used for verification and alignment of videographic and behavioral data (see above). The primary outcome measures were wheel-derived running speed and DeepLabCut-derived^25^ paw kinematics during air-on versus air-off epochs as these measures provide the most direct behavioral readout of locomotion and limb movement on the running wheel. Secondary outcome measures included optical-flow–based motion metrics, facial motion measures, and eye-region area.

Video data acquired from bottorm-view camera was analyzed using DeepLabCut, a markerless pose estimation framework based on deep learning ^25,26^. For this camera, the positions of the forepaws and hind paws were tracked to quantify limb kinematics during locomotion. Video preprocessing, pose estimation, and post-processing were performed using a combination of DeepLabCut utilities and custom MATLAB and Python scripts. Kinematic and pupil-derived measures were synchronized with encoder and stimulus signals using shared timing markers, enabling trial-by-trial alignment of locomotion, facial dynamics, and eye/pupil-related variables. All analysis scripts are made publicly available to facilitate replication and reuse of the analysis pipeline.

#### Video preprocessing

To reduce storage requirements and accelerate downstream computer-vision processing, the paws and face videos were downsampled in spatial resolution prior to analysis. Videos were resized to 25% of their original width and height (i.e., 1/4 scaling in each dimension) using OpenCV with area-based interpolation (INTER_AREA), which is well-suited for high-quality downscaling. The pupil video was recorded at a smaller native resolution and was not resized. Resized videos were saved alongside the originals with a “_reduced” suffix to preserve provenance.

#### ROI selection and metadata

For each video modality (paws, face, pupil), a rectangular region of interest (ROI) was selected manually on the first frame using an OpenCV interactive ROI tool. The ROI coordinates (x, y, width, height) and the source video path were saved to a sidecar JSON file (e.g., “*_roi.json”). This ensured that the same ROI definition could be reused for reprocessing, parameter tuning, and reproducible analysis across sessions.

#### Optical flow and motion-energy extraction

To obtain a fast, model-free characterization of movement dynamics without requiring pose-estimation networks, we computed dense optical flow within each ROI using the Dual TV-L1 method implemented in OpenCV (opencv-contrib). For each frame pair, the video was cropped to the ROI, converted to grayscale, and optical flow was estimated between successive frames. As summary motion features, we computed the mean horizontal and vertical flow components (avg_u, avg_v) within the ROI. In parallel, we computed a complementary motion-energy measure as the mean absolute frame-to-frame intensity difference within the ROI. For each modality, outputs were written to a CSV file containing frame index, timestamp (ms; from the video container), average flow components, and motion-energy values (e.g., “*_OF.csv”). Processing was performed offline.

#### LED-based synchronization of video streams

To synchronize independently acquired video streams with the air-delivery control signal, a red LED was driven by the same digital command used to actuate the air solenoid. The LED was positioned within the field of view of each behavioral camera (paws, face, and pupil). LED intensity was extracted from a small region of interest in each video and thresholded to generate a binarized air-state signal. Air-on and air-off events were identified from rising and falling transitions of this signal and used to align video-derived stimulus timing with the air-control signal recorded by the data acquisition (DAQ) system (5 kHz sampling).

### Statistical Analyses

Statistical analyses were performed using MATLAB (MathWorks). For the initial experiments presented here for the representative animal, trial-level analyses were performed. For each trial, behavioral measures such as running speed were quantified separately during air-on and the immediately preceding air-off epochs (phases). Trial-averaged values were then compared between phases using paired, two-tailed t-tests as well as Wilcoxon signed-rank test. In addition, distributions of speed values during air-on and air-off periods were compared using the Kolmogorov–Smirnov test.

For the additional cohort (N = 3 mice), cross-animal consistency was assessed using repeated-measures analysis of variance (RM-ANOVA), with air phase (air-on vs. air-off) treated as a within-subjects factor. Statistical significance was assessed at a threshold of p < 0.05. Data are reported as mean ± standard deviation for the trial-level analysis and mean ± standard error of the mean for animal-level analyses. Formal assessments of statistical assumptions (e.g., normality) were not used as strict inclusion criteria; therefore, both parametric and nonparametric results are reported where appropriate.

## Results

### Air-induced locomotion produces reliable trial structure on a transparent wheel

To determine whether air-stream–induced locomotion generates reliable and repeatable trial structure on a transparent running wheel, we first examined the relationship between air delivery and locomotor behavior in a head-fixed mouse. Running behavior during air-on epochs was quantitatively compared with behavior during air-off epochs within the same recording sessions. Continuous wheel-derived speed measurements revealed robust, stimulus-locked increases in running during air application, followed by decreases in speed upon air offset (Fig. 2A). Over an ∼11-minute recording session for the representative animal, the animal completed 34 trials in which air onset consistently triggered locomotion and air offset marked transitions to lower-speed states. In each trial, air delivery was maintained until the animal completed a fixed running distance of 25 cm, thereby defining discrete air-on and air-off epochs. To quantify the separation between locomotor states, we compared the distributions of running speed during air-on and air-off periods. Cumulative speed distributions showed a pronounced rightward shift during air-on epochs relative to air-off epochs (Fig. 2B), indicating robust modulation of locomotion by the air stimulus (Kolmogorov–Smirnov test, ***p < 0.001).

Next, we aligned running speed to stimulus transitions to assess temporal dynamics across trials. Following air onset, running speed increased rapidly and reached a stable plateau within approximately 1–2 seconds (Fig. 2C). Conversely, air offset was followed by a gradual decay in running speed, reflecting controlled disengagement from locomotion rather than abrupt stopping (Fig. 2D). These aligned averages demonstrate consistent temporal structure across trials despite natural variability in instantaneous speed.

Finally, to assess trial-to-trial reliability, mean running speed was computed for each air-on trial and compared with the immediately preceding air-off period. Across 34 trials, average speed during air-on epochs (mean ± SEM: 5.09 ± 0.18 cm/s) was significantly higher than during air-off epochs (2.93 ± 0.28 cm/s) (paired t-test, ***p < 0.001; Fig. 2E). To evaluate whether these effects generalize across animals, we repeated the analysis in a separate cohort of three mice. Consistent with the representative example, cumulative speed distributions again showed a clear separation between air-on and air-off states (Fig. 2F). Trial-aligned averages demonstrated rapid increases in running speed following air onset and decreases following air offset (Fig. 2G–H). Across animals, mean running speed during air-on epochs (mean ± SEM: 4.54 ± 0.52 cm/s) remained significantly higher than during air-off periods (0.66 ± 0.20 cm/s) (paired t-test, p < 0.01; Fig. 2I), confirming that the AIR paradigm reliably produces stimulus-locked locomotion across animals. These results demonstrate that air-induced locomotion on a transparent wheel produces reliable, repeatable trial structure suitable for quantitative behavioral analyses.

### Air-induced locomotion produces reliable paw kinematics on a transparent wheel

To determine whether air-stream–induced locomotion yields reliable and quantifiable limb kinematics on a transparent wheel, we first examined whole-body and paw-specific movement using complementary video-based analyses. Optical-flow and motion-energy metrics extracted from a paw-view video revealed robust, stimulus-locked increases in movement during air application (Fig. 4). Paw optical-flow–derived speed and motion energy increased rapidly following air onset and decreased following air offset, producing well-defined air-on and air-off epochs across repeated trials (Fig. 4B–D). Trial-averaged analyses aligned to air onset and offset showed consistent temporal structure, with a rapid rise in paw movement following air onset and a gradual decay following air termination (Fig. 4C,D).

Across trials (n = 34), mean paw speed during air-on epochs was significantly higher than during air-off epochs (paired t-test, p < 0.001; Fig. 4E), as was motion energy (p < 0.001; Fig. 4F). These results demonstrate that air delivery reliably evokes reproducible paw and body movement on the transparent wheel, providing a robust behavioral readout that is temporally aligned to the stimulus. To assess whether these kinematic patterns generalize across animals, the analysis was repeated for the separate cohort of three mice. Trial-aligned averages again showed rapid increases in optical-flow speeds following air onset and decreases following air offset (Fig. 3G,H). Across animals, speed remained higher during air-on epochs compared with air-off periods (Fig. 3I) while motion energy showed a similar trend (Fig. 3J).

**Figure 3.**
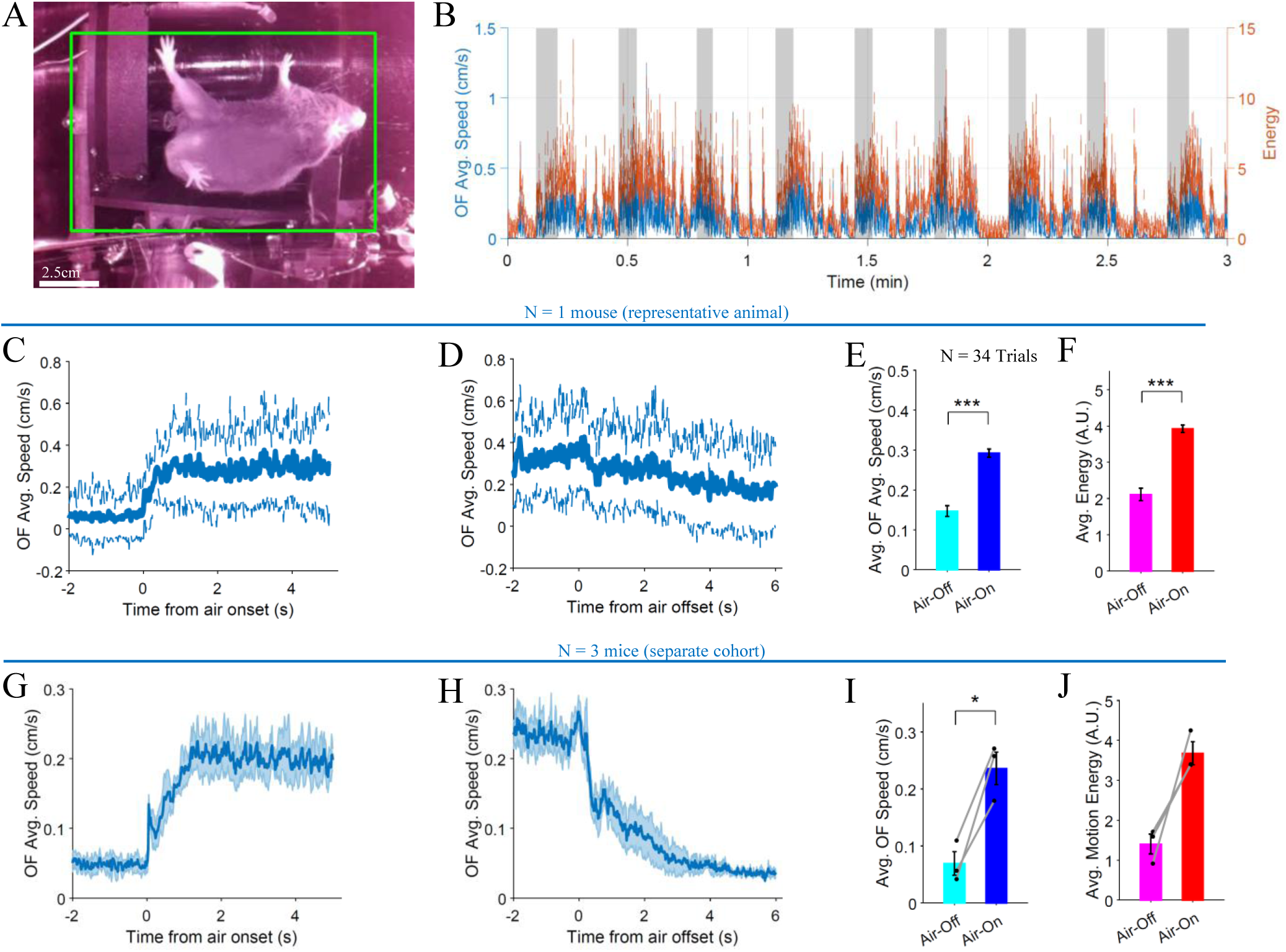
Whole-body kinematics during air-induced locomotion. **(A)** Representative paw video frame showing the region of interest (ROI; green box) used for optical-flow (OF) and motion-energy extraction. Example tracked landmarks are overlaid to illustrate paw position within the ROI (scale bar, 5 cm). **(B)** Time series of paw optical-flow–derived average speed (blue, left axis) and motion energy (orange, right axis) over a representative recording segment. Shaded gray regions denote air-on epochs identified from the LED-based air signal. **(C)** Trial-averaged OF speed aligned to air onset (t = 0 s). Solid line indicates the mean across trials; dashed lines indicate ± SD. **(D)** Trial-averaged OF speed aligned to air offset (t = 0 s), plotted as in (C). **(E)** Mean paw OF speed during air-on versus air-off epochs, averaged across trials (N = 34). Bars show mean ± SEM; ***p < 0.001 (paired t-test). **(F)** Mean paw motion energy during air-on versus air-off epochs (n = 34), plotted as in (E). **(G – J)** Same as C-F respectively but for a separate cohort of N = 3 mice. Lines and bar graphs represent mean over trials and animals and shaded regions represent the standard error of the mean across animals. Individual dots in (I) and (J) indicate mean values for individual animals.

To further resolve limb-specific kinematics, we applied DeepLabCut-based pose estimation to track individual body points, including all four paws, tail base, and nose (Fig. 4). Tracking confidence was high across all body parts, with the majority of frames exceeding a likelihood threshold of 0.9 and sharply peaked likelihood distributions (Fig. 4B, representative animal), indicating stable and reliable pose estimation throughout the session. Time-resolved trajectories of tracked points revealed structured, stimulus-locked changes in both horizontal and vertical paw positions during air-on epochs, consistent with coordinated locomotor behavior rather than sporadic or noisy movement.

**Figure 4.**
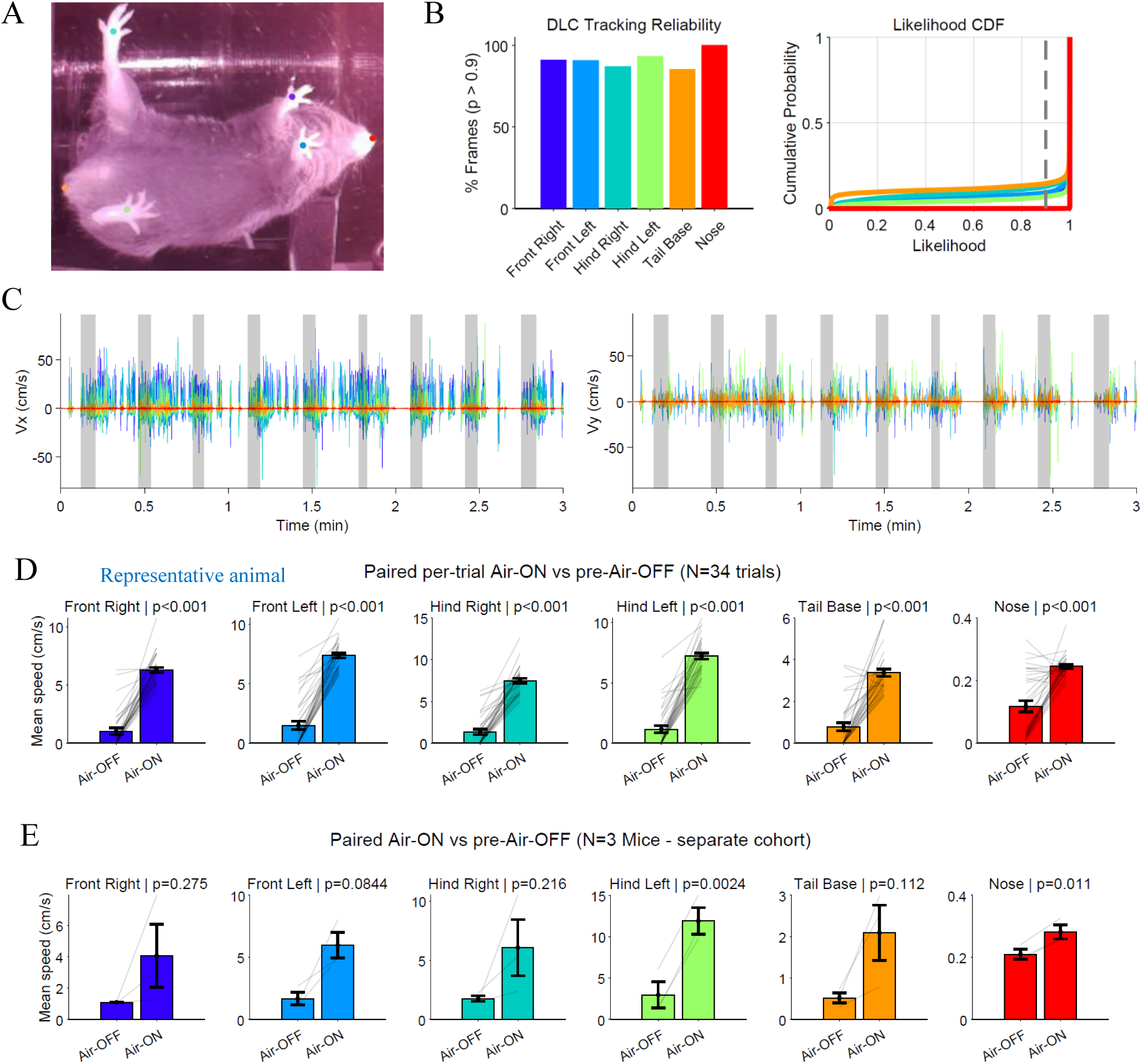
DeepLabCut-based paw and body kinematics during air-induced locomotion. **(A)** Example frame from the paw-view video showing DeepLabCut tracking of forelimbs, hindlimbs, tail base, and nose. Colored markers indicate tracked body points used for kinematic analysis. **(B)** Tracking reliability across body parts. Left: percentage of frames with likelihood > 0.9 for each tracked point. Right: cumulative distribution functions (CDFs) of likelihood values, demonstrating consistently high confidence across body parts. **(C)** Horizontal (X; left) and vertical (Y; right) speeds of all tracked body points over time. Shaded regions indicate air-on epochs, illustrating structured, stimulus-locked changes in paw and body position during air-induced locomotion. **(D)** Trial-wise comparison of mean paw and body-point speed during air-on versus pre-air-off periods (N = 34 trials) for the representative animal. Each gray line represents a single trial (paired comparison), bars show mean ± SEM. All tracked body points exhibited significantly higher movement speeds during air-on epochs (paired t-tests, all p < 0.001). **(E)** Same as D but for a separate cohort of N = 3 mice. Bar graphs represent mean over trials and animals with the standard error of the mean across animals. Individual dots indicate mean values for individual animals.

Quantifying per-trial movement speeds for each tracked body point showed that all paws, as well as the tail base and nose, exhibited significantly higher speeds during air-on periods compared to pre-air-off intervals (Fig. 4C and Fig. 4D, paired t-tests, all p < 0.001) with similar trends observed across animals in a separate cohort (Fig. 4E). The consistent increase across multiple body points confirms that air-induced locomotion on the transparent wheel engages coordinated whole-body movement rather than isolated limb motion. These analyses demonstrate that air-stream stimulation produces reliable, repeatable, and quantifiable paw and body kinematics in head-fixed mice running on a transparent wheel..

### Facial and Eye dynamics during air-induced locomotion

To assess stimulus-locked facial movements and autonomic correlates during air-induced locomotion, we analyzed front-facial and side-facial video streams acquired simultaneously with paw kinematics (Figs. 5 and 6). Optical-flow (OF) and motion-energy measures were extracted from predefined facial regions of interest to quantify gross facial motion, while DeepLabCut (DLC) was used to track discrete facial landmarks to evaluate tracking reliability and movement consistency.

**Figure 5.**
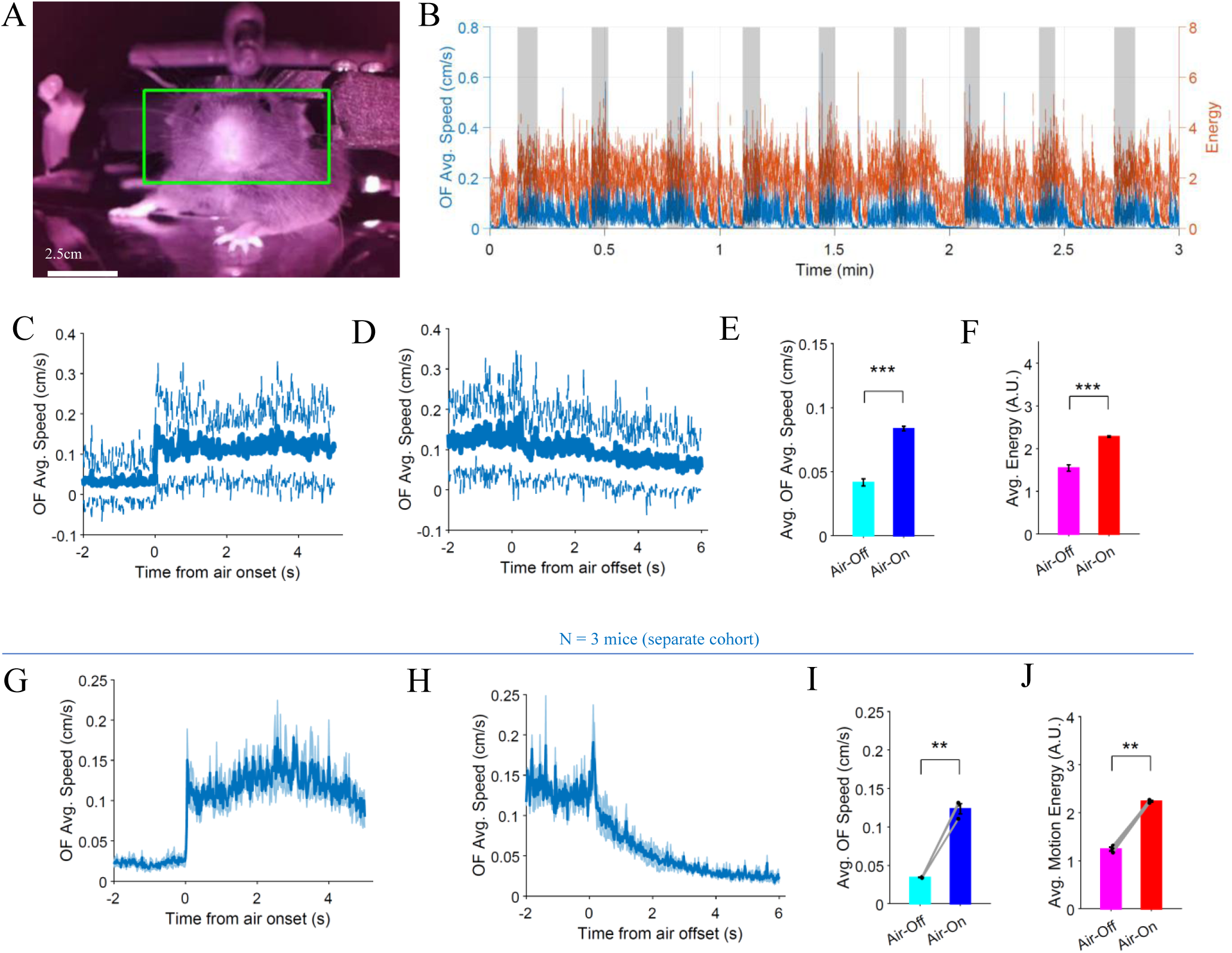
Front-facial kinematics during air-induced locomotion. **(A)** Representative front-facial video frame illustrating the region of interest (ROI; green box) used for optical-flow (OF) and motion-energy extraction (scale bar, 2.5 cm). **(B)** Time series of OF-derived average facial motion speed (blue, left axis) and motion energy (orange, right axis) during a representative recording segment. Shaded gray regions indicate air-on epochs identified from the LED-based air signal. **(C)** Trial-averaged facial OF speed aligned to air onset (t = 0 s). Solid line indicates the mean across trials; dashed lines indicate ± SD. **(D)** Trial-averaged facial OF speed aligned to air offset (t = 0 s), plotted as in (C). **(E)** Mean facial OF speed during air-on versus air-off epochs, averaged across trials. Bars show mean ± SEM; ***p < 0.001 (paired t-test). **(F)** Mean facial motion energy during air-on versus air-off epochs, plotted as in (E). **(G – J)** Same as C-F respectively but for a separate cohort of N = 3 mice. Lines and bar graphs represent mean over trials and animals and the shaded regions represent the standard error of the mean across animals. Individual dots in (K) and (L) indicate mean values for individual animals.

**Figure 6.**
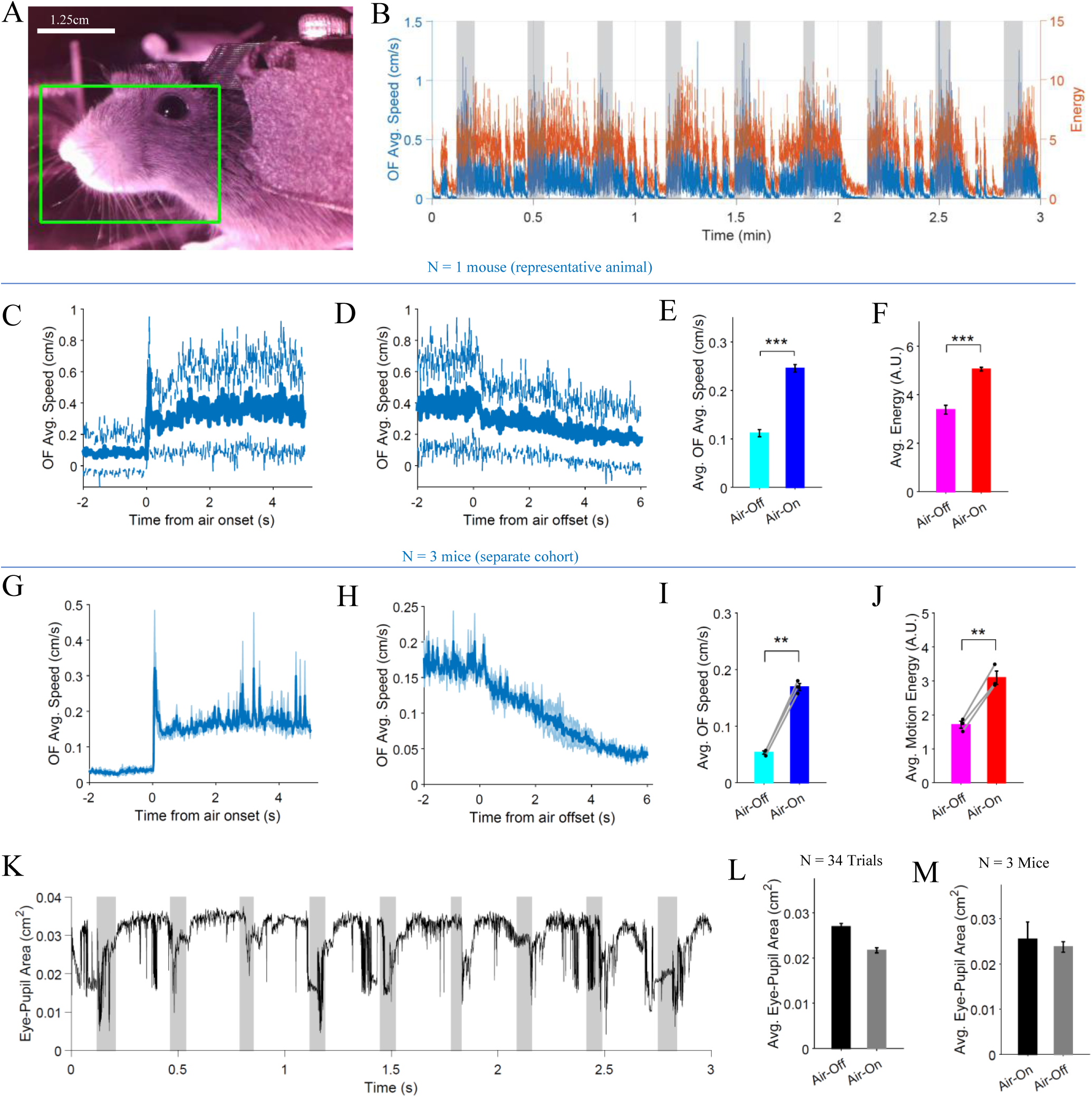
Side-facial and eye dynamics during air-induced locomotion. **(A)** Representative side-facial video frame showing the region of interest (ROI; green box) used for optical-flow (OF) and motion-energy extraction from the facial and periocular region (scale bar, 1.25 cm). **(B)** Time series of OF-derived average facial motion speed (blue, left axis) and motion energy (orange, right axis) during a representative recording segment. Shaded gray regions indicate air-on epochs identified from the LED-based air signal. **(C)** Trial-averaged facial OF speed aligned to air onset (t = 0 s). Solid line denotes the mean across trials; dashed lines indicate ± SD. **(D)** Trial-averaged facial OF speed aligned to air offset (t = 0 s), plotted as in (C). **(E)** Mean facial OF speed during air-on versus air-off epochs, averaged across trials. Bars show mean ± SEM; **p < 0.01, ***p < 0.001 (paired t-test). **(F)** Mean facial motion energy during air-on versus air-off epochs, plotted as in (E). **(G– J)** Same as C-F respectively but for a separate cohort of N = 3 mice. Lines and bar graphs represent mean over trials and animals and shaded regions represent the standard error of the mean across animals. Individual dots in (K) and (L) indicate mean values for individual animals. **(K)** Time series of eye-region area extracted from side-facial video. **(L)** Mean eye-region area during air-on versus air-off epochs. **(M)** Same as (L) but from an experiment with a separate cohort of N = 3 mice.

In the front-facial view, air delivery produced a clear and reproducible increase in facial motion. Time-resolved OF-derived facial speed and motion energy closely tracked air-on epochs identified from the LED signal (Fig. 5B). Trial-aligned analyses revealed a rapid increase in facial motion following air onset (Fig. 5C) and a gradual decay following air offset (Fig. 5D). Across trials, mean facial OF speed and motion energy were significantly higher during air-on compared to air-off epochs (paired t-tests, ***p < 0.001; Fig. 5E–F), indicating robust stimulus-locked facial activation during locomotion. A similar temporal structure and increase in facial motion during air-on epochs was observed in a separate cohort of three mice, with consistent trial-averaged dynamics and elevated OF speed and motion energy during air-on periods (Fig. 5G-J).

Side-facial recordings provided complementary measurements of facial motion and eye dynamics (Fig. 6). As in the front-facial view, OF-derived facial speed and motion energy increased reliably during air-on epochs (Fig. 6B–F), confirming that air delivery evokes coordinated facial movement across viewing angles. The same stimulus-locked increases in facial motion were also observed in a separate cohort of three mice (Fig. 6G–J), demonstrating that these effects generalize across animals.

In contrast, analysis of the eye region revealed a modest trend toward reduced eye or pupil area during air on epochs, although this effect did not reach statistical significance (Fig. 6K-M for representative animal and the separate cohort of N = 3 mice). Experiments were conducted in darkness using infrared illumination, resulting in a maximally dilated pupil throughout the session. Under these conditions, neither air delivery nor the red LED used to indicate air onset produced detectable changes in eye or pupil area, indicating that any stimulus-associated modulation of pupil size was minimal under these recording conditions.

These results demonstrate that air-induced locomotion is accompanied by robust, stimulus-locked facial movements that can be quantified using optical-flow–based measures. At the same time, the absence of pupil modulation under dark conditions establishes a useful baseline for interpreting future autonomic measurements obtained under different lighting or task contingencies.

### LED-based synchronization enables alignment of air stimulus across video streams

LED-based synchronization enabled alignment of the air stimulus across independently acquired video streams. A red LED driven by the same control signal as the air solenoid was positioned within the field of view of each camera (paws, face, and pupil), allowing air-on and air-off events to be detected directly from the video data. LED intensity signals were extracted independently from each video stream and converted into binarized air-state vectors using a data-driven single-threshold detection approach. Air-on and air-off epochs were then identified from rising and falling transitions in the binarized LED signal, providing independent estimates of air delivery timing for each video modality. Visual inspection of LED-aligned frame montages confirmed reliable detection of air onset across all three camera views, with frame-by-frame transitions occurring within the expected temporal resolution imposed by the video sampling rate (set to 60 Hz; Fig. 7A). The effective frame rates measured from frame counts and the whole recording duration obtained from DAQ were 60.53 Hz for the paws camera, 60.24 Hz for the face camera, and 62.41 Hz for the pupil camera, reflecting small deviations from the nominal acquisition rate due to independent camera clocks. When plotted across the full recording session, the binarized air-state signals derived from the paws, face, and pupil videos closely reproduced the trial structure of the air stimulus recorded by the DAQ system (Fig. 7B). Small offsets in absolute timing were occasionally observed between streams, reflecting independent camera clocks rather than loss of stimulus information.

**Figure 7.**
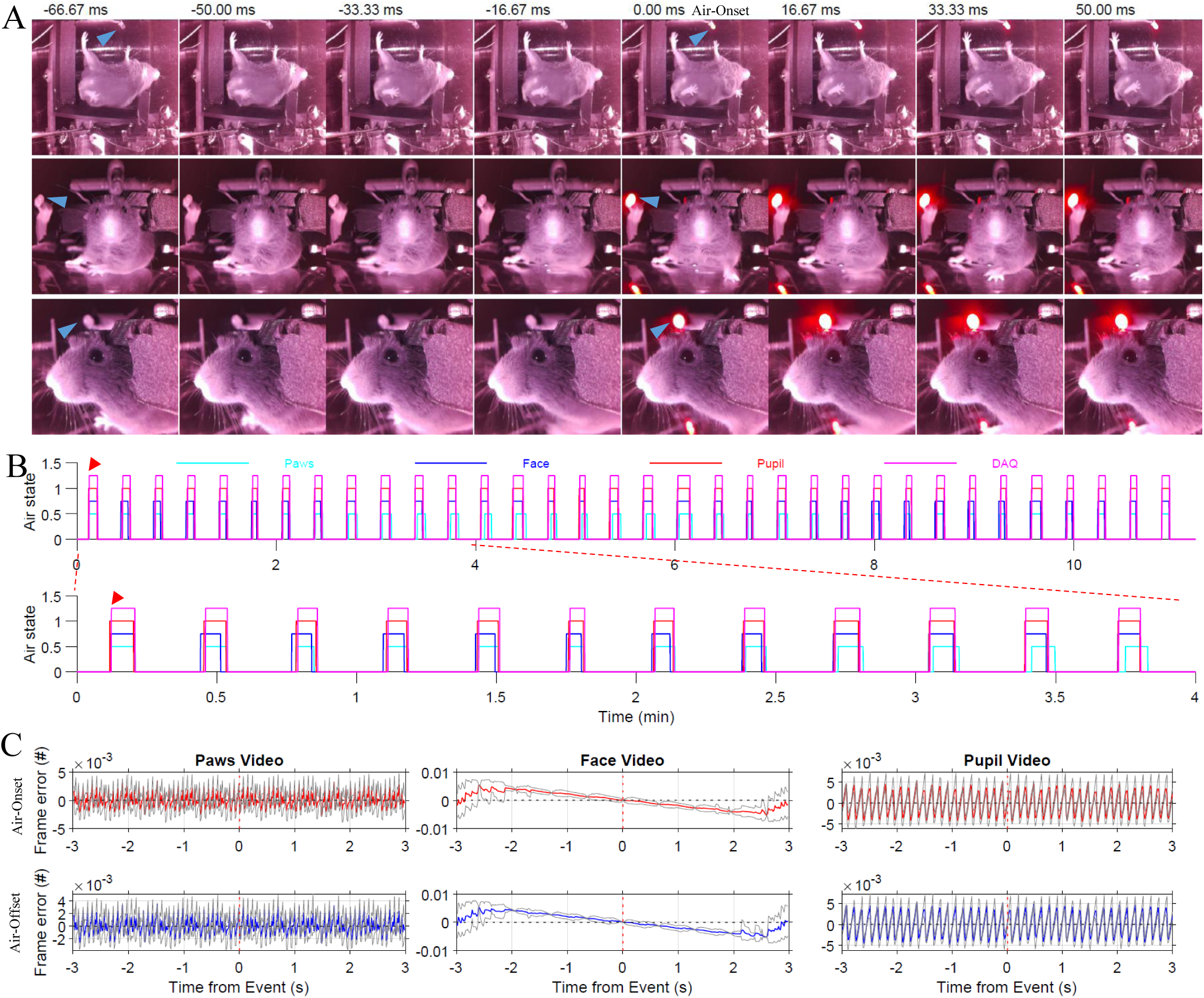
LED-based synchronization enables reliable alignment of air stimulus across behavioral and video modalities. **(A)** Representative montages from the three simultaneously acquired video streams (top: paws/bottom view; middle: face/front view; bottom: pupil/side view) aligned to the first detected air-onset event (indicated with red arrow head in panel B) in each video using the LED signal visible in the frame. Frames are shown from −66.7 ms to +50.0 ms relative to air onset (60 Hz sampling). Cyan arrowheads indicate the LED location within each field of view, demonstrating consistent visual detection of air onset across cameras despite differences in perspective. **(B)** Binarized air-on state extracted independently from LED intensity signals in the paws (cyan), face (blue), and pupil (red) videos, plotted over the full recording session with DAQ air signal (magenta). Although small offsets in absolute timing are visible across modalities due to independent acquisition clocks, trial structure and air-on epochs are preserved across all streams. **(C)** Quantification of synchronization accuracy between video streams and the DAQ system. Frame timing divergence was computed after anchoring signals at air-on (top row) and air-off (bottom row) events. Error is expressed in video frames. Across cameras, synchronization errors remained well below one frame (<0.01 frames) within ±3 s of the event, corresponding to sub-millisecond temporal alignment between independently acquired video streams and the DAQ system.

To quantify synchronization accuracy, LED-derived air-on and air-off events detected in each video stream were compared to the DAQ-recorded air-control signal (5 kHz sampling), which served as the temporal reference. After anchoring signals at air-on and air-off events, divergence between video and DAQ time axes was evaluated within ±3 s windows. Across cameras, synchronization errors remained well below one video frame (<0.01 frames), corresponding to sub-millisecond timing differences relative to the DAQ signal (Fig. 7C). These results indicate that the small differences in achieved frame rates across cameras do not affect stimulus alignment. These small deviations reflect minor differences in camera clock rates and do not accumulate sufficiently to affect event alignment within the analyzed time window.

## Discussion

The primary goal of this study was to develop and validate a modular, behavioral platform that enables locomotion on demand in head-fixed mice while providing synchronized, multi-view readouts of behavior. A central technical innovation of the platform is the use of a transparent running wheel (with the air-induced running task), which permits unobstructed videographic access to the ventral body surface and limbs during locomotion. This design enables simultaneous measurement of whole-body kinematics, paw dynamics, facial motion, and eye/pupil-related signals within a single experimental configuration, without requiring repositioning of the animal or cameras. Additional methodological novelties include stimulus-induced control of locomotion using air delivery, LED-based temporal synchronization across independent video streams, and the integration of optical-flow–based motion metrics with pose-estimation approaches. Importantly, the results presented here are intended as a technical proof of principle rather than a population-level characterization of behavioral phenomena. Accordingly, initial system validation and detailed behavioral characterization were performed using a representative, well-trained animal, allowing focused evaluation of system feasibility, reliability, temporal precision, and multi-view synchronization without confounds introduced by inter-animal variability. To assess cross-animal feasibility and reproducibility of the AIR paradigm with the transparent wheel, we subsequently incorporated an additional cohort of three animals (total N = 4). By establishing stable and reproducible performance both at the representative-subject level and across animals, the present work provides a robust foundation for future studies employing larger cohorts and simultaneous neural recordings.

A key conceptual contribution of this platform is the integration of stimulus-induced locomotion with simultaneous, multi-view behavioral readout under head fixation in contrast to previously reported transparent surface locomotion systems ^14–16^. The transparent wheel provides direct ventral imaging access to the limbs, enabling visualization and quantification of paw kinematics that are difficult to obtain with opaque or belt-based running systems. Ventral-view recordings show cyclic stepping patterns during locomotion (see raw behavioral videos provided on OSF), although the smooth transparent surface differs in frictional properties from belt-based treadmills and may influence fine kinematic details. During active locomotion, animals exhibit coordinated stepping movements consistent with natural wheel walking or running. Animals may also occasionally orient sideways on the wheel during locomotion or stationary periods, which can produce outward positioning of the paws relative to the body. In addition, the ventral viewing geometry may visually exaggerate this appearance compared with lateral perspectives. When combined with synchronized facial videography, this configuration allows concurrent assessment of locomotor output, facial dynamics, and eye/pupil-related signals within the same trials, using a single unified stimulus paradigm.

An important advantage of the AIR task is that it provides a rapid, repeatable, and minimally structured means of inducing locomotion without reliance on reward learning, deprivation, or extended shaping procedures ^24,27–29^. Air-induced locomotion exploits an innate sensorimotor response that can be established reliably within a few sessions, producing behavior with well-defined onset and offset transitions. In this study task structure was acquired within relatively short training periods (2-14 days) and is in agreement with previous studies with related treadmill-based implementation of the AIR paradigm where training times were 3-5 days ^18^. This structure enables precise segmentation of immobility-to-locomotion and locomotion-to-immobility epochs, which is particularly advantageous for studying state transitions, motor initiation, and context-dependent modulation of neural and behavioral variables ^30^. Moreover, because air termination is behavior-dependent rather than fixed in duration, the paradigm may couple stimulus delivery to performance, allowing flexible manipulation of task demands (e.g., target distance or effort) without altering the underlying control logic. These features position the AIR task as a complementary alternative to reward-based locomotion assays, particularly for experiments focused on temporal dynamics and multimodal state changes.

A central design objective of the AIR task was to achieve precise temporal alignment between stimulus delivery and multi-view behavioral measurements acquired across heterogeneous hardware and sampling rates. By combining Arduino-controlled air delivery with LED-based visual markers embedded directly within each video stream, the system provides a robust and redundant synchronization strategy that does not rely on shared clocks or post hoc assumptions about timing. Quantitative comparison of air-on and air-off durations extracted from LED-derived signals against DAQ-recorded air-control signals demonstrated high agreement across video modalities, with deviations largely attributable to the temporal discretization imposed by video frame rates. Importantly, these discrepancies were small relative to the duration of air-on and air-off epochs and do not constrain within-epoch event-related analyses of locomotion, kinematics, or facial dynamics. This synchronization approach ensures reliable stimulus alignment even when video recordings operate at different sampling frequencies, a common challenge in multi-view experimental designs. This framework also enables precise temporal registration of neural recordings with video-based behavioral measurements in future multimodal experiments.

The platform described here is readily extensible to a wide range of experimental applications. Future studies will incorporate larger cohorts to enable population-level inference and to assess inter-animal variability in locomotor, facial, and autonomic responses. Critically, the system is designed to integrate seamlessly with simultaneous neural recordings, including two-photon calcium imaging, electrophysiology, and fiber photometry, enabling direct investigation of how neural activity across cortical and subcortical circuits relates to stimulus-evoked locomotion and behavioral state transitions. The modular design further allows straightforward adaptation of task parameters, such as varying air intensity, target distance, or sensory context, to probe graded motor effort, motivation, and sensorimotor integration. Beyond basic motor control, this framework is well suited for studying early alterations in movement, arousal, and coordination in disease models, including aging and neurodegenerative disorders, where subtle behavioral changes may precede overt impairments.

## Limitations

Several limitations of the current platform should be noted. First, the cameras used in this implementation operate on independent internal clocks and are not hardware-synchronized. Although LED-based event detection enables reliable alignment of behavioral epochs across video streams, precise frame-by-frame synchronization across cameras is not guaranteed. Future implementations using hardware-triggered cameras could enable true frame-locked acquisition and facilitate applications requiring cross-camera 3D reconstruction or integration with high temporal-resolution neural recordings such as electrophysiology ^31,32^. Second, although facial and whisker motion can be detected during air delivery, a portion of this movement may reflect direct mechanical effects of the air stream rather than exclusively self-generated behavior. Whiskers, in particular, are lightweight structures susceptible to passive deflection. This limitation could be mitigated in future implementations by incorporating a simple air shield or baffle positioned to protect the face and whisker pad while preserving airflow to the body. Third, while the platform includes a pupil camera capable of supporting pupillometry, pupil dynamics were not systematically manipulated or quantified in the present study. Future work incorporating controlled illumination or stimulus-driven pupil modulation will allow this capability to be fully validated ^22,26^. Fourth, although the platform enables multi-angle videography of body and facial kinematics, whisker tracking was not achieved in the present dataset. Optimized whisker tracking approaches may allow future studies to incorporate whisker kinematics ^12^. Finally, while the system reliably evokes locomotion, systematic characterization of airflow parameters and their relationship to locomotor speed was beyond the scope of this initial validation study.

## Supporting information

AWB_Assembly_and_Operations_Guide

Arduino Code

Matlab Code

Raspberry Pi Code

3D printing files

## Resource Availability

A complete list of materials, detailed design schematics with assembly guide, and 3D-printable files for all custom components used in this setup are provided in the Supplementary Materials. All designs, hardware specifications, and associated software are made available to facilitate replication and adaptation of the system by other laboratories.

### Lead contact

Further information and requests for resources and reagents should be directed to and will be fulfilled by the lead contact, Samsoon Inayat.

### Materials availability

This study did not generate new unique reagents.

### Data and code availability

All the raw data generated and analyzed during the current study are publicly available on the Open Science Framework (OSF) repository, [https://osf.io/k9j3b/overview]. All custom analysis code used in this study is publicly available on GitHub at [https://github.com/neuromomentumlab/AIR_Wheel_Methods].

### Protocol registration

No formal protocol registration was performed prior to this study

## Author Contributions

All authors contributed to the design of the study. P.P. and S.I. led the design and integration of the behavioral apparatus. P.P. designed and fabricated the 3D-printed components. T.M.G. participated in system integration and performed the animal surgeries. T.M.G., P.P., and S.I. conducted the behavioral experiments. P.P. and S.I. developed the Arduino, Raspberry Pi, and MATLAB code for stimulus control, data acquisition, and synchronization. P.P. and S.I. implemented the synchronization and alignment analyses. P.P. and T.M.G. performed the DeepLabCut-based pose estimation analyses. S.I. performed the optical-flow–based motion analyses and wrote the first draft of the manuscript. All authors reviewed and approved the final manuscript.

## Acknowledgements

We thank the animal care staff at the University of Nevada, Las Vegas for their support and assistance with animal husbandry. We also thank Dr. John Troy, Professor Emeritus, Northwestern University, Evanston, Illinois for donating equipment used for this study including CNC machine tools and the NI-USB 6259 data acquisition card.

## Funding

This work was supported by an Interdisciplinary Neuroscience Graduate Assistantship to Pratik Paranjape, Department of Psychology Graduate Assistantship to Tahoura Mohammadi Ghohaki, and internal UNLV funding to Samsoon Inayat.

## Declaration of Interests

The authors declare that an invention disclosure related to the behavioral setup described in this manuscript has been filed with the University of Nevada, Las Vegas.

## Notes

### Summary of Updates

Addition of data and analyses from 3 more animals

## References

1. Becker, M. I., Calame, D. J., Wrobel, J. & Person, A. L. Online control of reach accuracy in mice. Journal of Neurophysiology 124, 1637–1655 (2020).

2. Yue, Y. et al. Motor training improves coordination and anxiety in symptomatic Mecp2-null mice despite impaired functional connectivity within the motor circuit. SCIENCE ADVANCES (2021).

3. Luong, T. N., Carlisle, H. J., Southwell, A. & Patterson, P. H. Assessment of Motor Balance and Coordination in Mice using the Balance Beam. JoVE 2376 (2011) doi:10.3791/2376.

4. Heindorf, M., Arber, S. & Keller, G. B. Mouse Motor Cortex Coordinates the Behavioral Response to Unpredicted Sensory Feedback. Neuron 99, 1040–1054.e5 (2018).

5. Nguyen, K. P., Sharma, A., Gil-Silva, M., Gittis, A. H. & Chase, S. M. Distinct Kinematic Adjustments over Multiple Timescales Accompany Locomotor Skill Development in Mice. Neuroscience 466, 260–272 (2021).

6. Vinck, M., Batista-Brito, R., Knoblich, U. & Cardin, J. A. Arousal and locomotion make distinct contributions to cortical activity patterns and visual encoding. Neuron 86, 740–754 (2015).

7. Sun, G. et al. Neural representation of self-initiated locomotion in the secondary motor cortex of mice across different environmental contexts. Commun Biol 8, 725 (2025).

8. Quarta, E. et al. Distributed and Localized Dynamics Emerge in the Mouse Neocortex during Reach-to-Grasp Behavior. J. Neurosci. 42, 777–788 (2022).

9. Currie, S. P. et al. Movement-specific signaling is differentially distributed across motor cortex layer 5 projection neuron classes. Cell Reports 39, 110801 (2022).

10. Carlsen, E. M. M., Nedergaard, M. & Rasmussen, R. N. Versatile treadmill system for measuring locomotion and neural activity in head-fixed mice. STAR Protocols 3, 101701 (2022).

11. Juczewski, K., Koussa, J. A., Kesner, A. J., Lee, J. O. & Lovinger, D. M. Stress and behavioral correlates in the head-fixed method: stress measurements, habituation dynamics, locomotion, and motor-skill learning in mice. Sci Rep 10, 12245 (2020).

12. Warren, R. A. et al. A rapid whisker-based decision underlying skilled locomotion in mice. eLife 10, e63596 (2021).

13. Krishnan, S. et al. A contextual fear conditioning paradigm in head-fixed mice exploring virtual reality. eLife 14, RP105422 (2025).

14. Machado, A. S., Darmohray, D. M., Fayad, J., Marques, H. G. & Carey, M. R. A quantitative framework for whole-body coordination reveals specific deficits in freely walking ataxic mice. eLife 4, e07892 (2015).

15. Darmohray, D. M., Jacobs, J. R., Marques, H. G. & Carey, M. R. Spatial and Temporal Locomotor Learning in Mouse Cerebellum. Neuron 102, 217–231.e4 (2019).

16. Albrecht, B., Schatz, A., Frei, K. & Winter, Y. KineWheel-DeepLabCut Automated Paw Annotation Using Alternating Stroboscopic UV and White Light Illumination. eNeuro 11, ENEURO.0304-23.2024 (2024).

17. Giovannucci, A. et al. Automated gesture tracking in head-fixed mice. Journal of Neuroscience Methods 300, 184–195 (2018).

18. Inayat, S., McAllister, B. B., Whishaw, I. Q. & Mohajerani, M. H. Hippocampal conjunctive and complementary CA1 populations relate sensory events to movement. iScience 26, 106481 (2023).

19. Inayat, S., McAllister, B. B., Whishaw, I. Q. & Mohajerani, M. H. Distinct Neural Signatures of Hippocampal Population Dynamics During Locomotion-in-Place. 2025.10.28.685240 Preprint at 10.1101/2025.10.28.685240 (2025).

20. Niell, C. M. & Stryker, M. P. Modulation of visual responses by behavioral state in mouse visual cortex. Neuron 65, 472–9 (2010).

21. Clack, N. G. et al. Automated Tracking of Whiskers in Videos of Head Fixed Rodents. PLoS Comput Biol 8, e1002591 (2012).

22. Bushnell, M., Umino, Y. & Solessio, E. A system to measure the pupil response to steady lights in freely behaving mice. Journal of Neuroscience Methods 273, 74–85 (2016).

23. Mao, D. et al. Hippocampus-dependent emergence of spatial sequence coding in retrosplenial cortex. Proc Natl Acad Sci U S A 115, 8015–8018 (2018).

24. Mao, D., Kandler, S., McNaughton, B. L. & Bonin, V. Sparse orthogonal population representation of spatial context in the retrosplenial cortex. Nat Commun 8, 243–243 (2017).

25. Mathis, A. et al. DeepLabCut: markerless pose estimation of user-defined body parts with deep learning. Nature neuroscience 21, 1281–1289 (2018).

26. Kobayashi, G. Pupil Dynamics-derived Sleep Stage Classification of a Head-fixed Mouse Using a Recurrent Neural Network.

27. Taxidis, J. et al. Differential Emergence and Stability of Sensory and Temporal Representations in Context-Specific Hippocampal Sequences. Neuron 108, 984–998 e9 (2020).

28. Itskov, P. M., Vinnik, E. & Diamond, M. E. Hippocampal representation of touch-guided behavior in rats: persistent and independent traces of stimulus and reward location. PLoS One 6, e16462 (2011).

29. Vedder, L. C., Miller, A. M. P., Harrison, M. B. & Smith, D. M. Retrosplenial Cortical Neurons Encode Navigational Cues, Trajectories and Reward Locations During Goal Directed Navigation. Cereb. Cortex 10.1093/cercor/bhw192 (2016) doi:10.1093/cercor/bhw192.

30. Arriaga, M. & Han, E. B. Dedicated Hippocampal Inhibitory Networks for Locomotion and Immobility. The Journal of neuroscience : the official journal of the Society for Neuroscience 37, 9222–9238 (2017).

31. Khilkevich, A. et al. Brain-wide dynamics linking sensation to action during decision-making. Nature 634, 890–900 (2024).

32. Stringer, C. et al. Spontaneous behaviors drive multidimensional, brainwide activity. Science 364, 255 (2019).

