## Supplementary material for "A TRANSPARENT WHEEL-BASED PLATFORM FOR LOCOMOTION-ON-DEMAND AND MULTI-VIEW BODY AND FACIAL KINEMATICS IN HEAD-FIXED MICE": AWB_Assembly_and_Operations_Guide

### AIR-Wheel Behavioral Platform

#### Assembly and Operations Guide

**Version:** 1.0
**Year:** 2026

Department of Psychology
University of Nevada, Las Vegas
Las Vegas, Nevada, USA

---

#### About this Guide

This document describes the **assembly, configuration, and operation** of the AIR-Wheel behavioral platform used for head-fixed locomotion experiments in mice.

The guide provides step-by-step instructions for constructing the mechanical apparatus, installing sensors and cameras, configuring the electronics and control systems, and running behavioral experiments. It also outlines the data acquisition pipeline and the processing workflow used to synchronize behavioral, video, and stimulus signals.

The AIR-Wheel system combines:

- Transparent running-wheel locomotion
- Air-induced stimulus control
- Multi-view behavioral videography
- Encoder-based locomotion measurement
- MATLAB-based data acquisition
- Raspberry Pi–based video recording

This modular design allows simultaneous measurement of locomotion, paw kinematics, facial dynamics, and eye/pupil signals in head-fixed mice while maintaining compatibility with neural imaging and electrophysiology systems.

#### Table of Contents


#### Assembly Guide for the AIR-Wheel Behavioral Platform

- Overview

Step 1. Base Plate Preparation

Step 2. Transparent Running Wheel Installation

Step 3. Encoder Wheel Assembly

Step 4. Head-Fixation Assembly

Step 5. Animal Casing

Step 6. Camera Mounting

Step 7. Air Delivery System

Step 8. Electrical Connections

---

#### System Operation and Data Acquisition Workflow

Step 1: Initialize the DAQ system

Step 2: Start Raspberry Pi camera scripts

Step 3: Begin MATLAB recording

Step 4: Behavioral recording

Step 5: End recording

Step 6: Data output

---

#### Data Organization and Loading Workflow

Raw Data Structure


---

#### Data Processing Workflow

##### Step 1: Define project directories

##### Step 2: Load experiment metadata

##### Step 3: Process behavioral signals

##### Step 4: Convert and process videos

##### Step 5: Load LED synchronization signals

##### Step 6: Load pupil region and signals

##### Step 7: Load video files for analysis

---

#### Behavioral Signal Processing (MATLAB)

##### Loading recorded behavioral data

##### Detecting stimulus transitions

##### Encoder signal processing

- Quadrature encoder channels
- Direct encoder count

##### Distance calculation

##### Running speed calculation

##### Speed smoothing

---

#### Video Processing and Frame-Rate Correction (MATLAB)

##### Processing Workflow

1. Identify recorded video files
2. Obtain the true recording duration
3. Count the number of frames in the video
4. Compute the effective frame rate
5. Convert the raw video file to MP4
6. Save video metadata

##### Final Output

### Assembly Guide for the AIR-Wheel Behavioral Platform

#### Overview

The AIR-Wheel apparatus consists of a base plate supporting a transparent running wheel, head-fixation assembly, multi-camera videography system, and air-delivery mechanism (Fig. AG-S0). The system is constructed using standard aluminum rods and clamps commonly used in optical and electrophysiology setups.
The steps below describe the recommended assembly procedure.

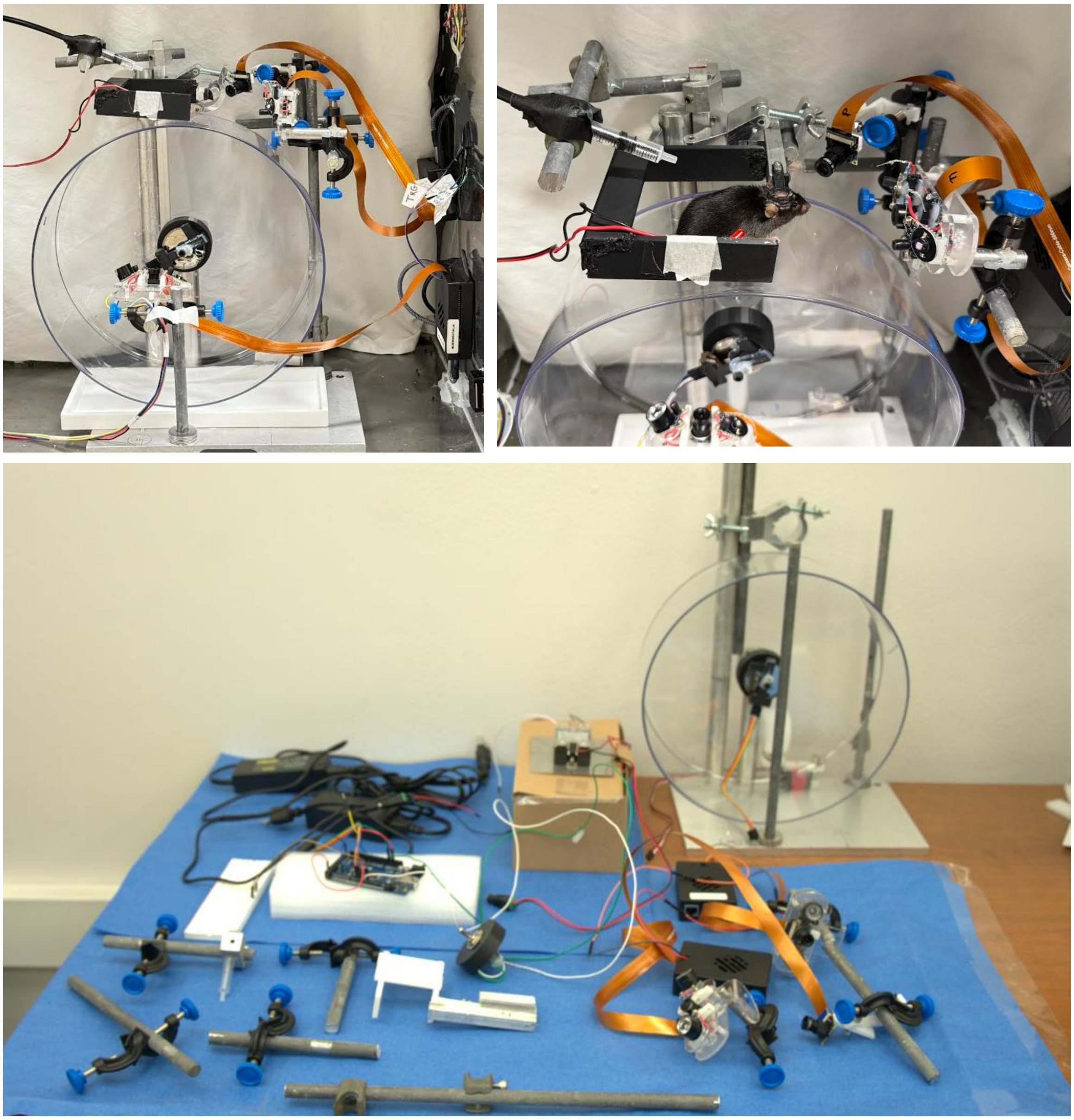

*Figure AG-S0. Overview of the AIR-Wheel behavioral platform and major system components. Photographs showing the complete AIR-Wheel experimental platform and its primary mechanical and electronic components used for behavioral experiments. The system includes the transparent running wheel mounted on an aluminum base plate with vertical support rods, a rotary encoder for measuring wheel rotation, head-fixation and animal casing components, and camera mounting hardware used for recording facial, pupil, and paw movements. Also shown are the control electronics, including the Arduino microcontroller, wiring for airflow stimulation and encoder signals, and associated power and communication connections used for synchronized data acquisition. This overview illustrates the modular design of the AIR-Wheel system prior to final assembly of the individual subsystems described in subsequent figures.*

#### Step 1. Base Plate Preparation

The apparatus is mounted on a rigid base plate (aluminum in our implementation).

Four vertical support rods are mounted on the base plate:

• Two 1-inch diameter circular aluminum rods
• Two ½-inch diameter circular aluminum rods

The two 1-inch rods serve distinct purposes:

1. The first 1-inch rod supports the transparent running wheel.
2. The second, taller 1-inch rod supports the head-fixation assembly using adjustable rod clamps.

The two ½-inch rods serve as auxiliary supports used for mounting cameras and other experimental components.

If the base plate does not contain pre-drilled mounting holes, mounting holes can be drilled to position the rods appropriately.
Alternatively, a commercial optical breadboard can be used where rods attach directly using threaded holes.

#### Step 2. Transparent Running Wheel Installation

The transparent running wheel is mounted on the first 1-inch aluminum rod.

Most commercially available transparent hamster wheels include a cylindrical mounting hub designed to fit onto a rod.

1. Slide the wheel's mounting hub onto the rod.
2. Align the wheel so that it rotates freely.
3. Tighten the locking screw provided with the wheel to secure it.

This configuration allows the wheel to rotate freely while maintaining mechanical stability.

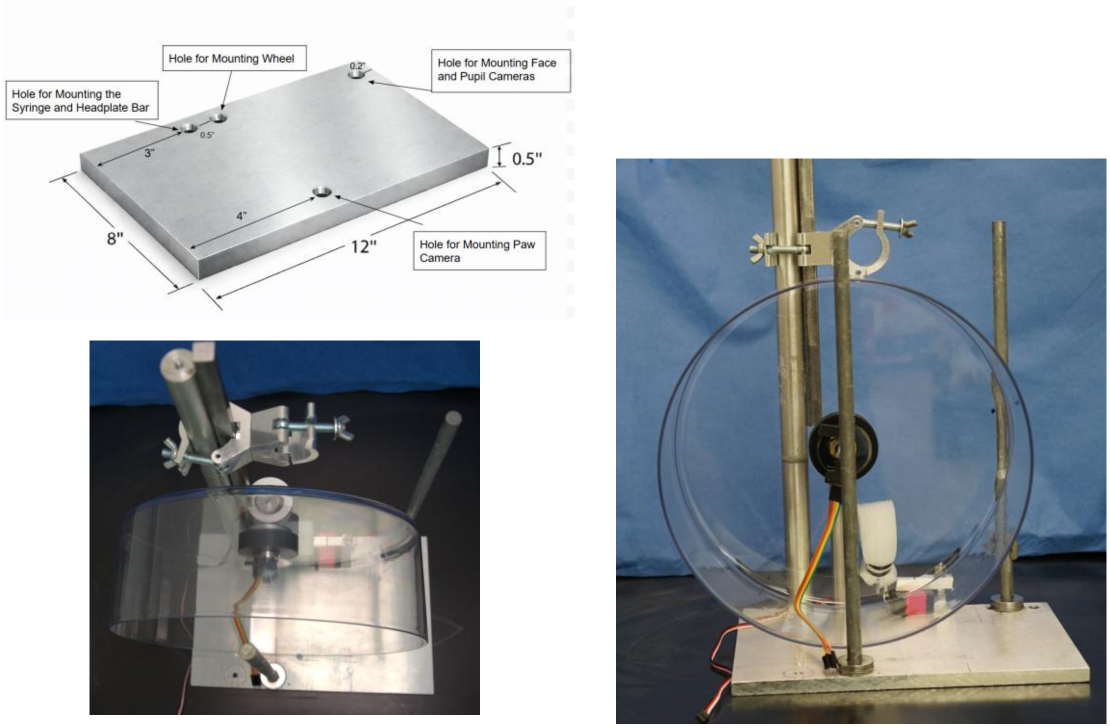


*Figure AG-S1. Base platform and transparent wheel assembly. The AIR-Wheel apparatus is constructed on a rigid aluminum base plate supporting multiple vertical metal rods used for mounting mechanical and imaging components. Two 1-inch diameter circular aluminum rods are mounted to the base plate. The first rod supports the transparent running wheel, which is mounted through the wheel hub and aligned to rotate freely. The second, taller rod supports the head-fixation assembly, which is attached using adjustable rod clamps at the top of the rod. An additional ½-inch diameter rod is positioned on the base plate to support auxiliary components such as camera mounts. The rotary encoder wheel assembly is positioned behind the transparent wheel and mechanically coupled to the wheel hub to measure wheel rotation. Left: Top view of the base plate showing the spatial arrangement of the rods, transparent wheel, and encoder assembly. Right: Front view of the platform illustrating the vertical alignment of the rods and the mounting position of the running wheel.*

#### Step 3. Encoder Wheel Assembly

To measure wheel rotation and locomotion speed, a rotary encoder is attached to the wheel using a custom 3D-printed cylindrical adapter and ball bearing.

Assembly principle:
• The outer bearing rotates with the running wheel.
• The inner bearing remains fixed and connects to the encoder transceiver.

Installation procedure:
1. Insert the 3D-printed cylindrical adapter onto the wheel's inside.
2. Attach the encoder wheel to the adapter.
3. Attach the encoder transceiver module to the fixed inner bearing structure.

Most transparent wheels include a threaded insert in the center hub. Remove the default screw and replace it with a screw used to attach the encoder assembly.

When the wheel rotates:
• The encoder wheel rotates with the outer bearing.
• The encoder sensor remains stationary.

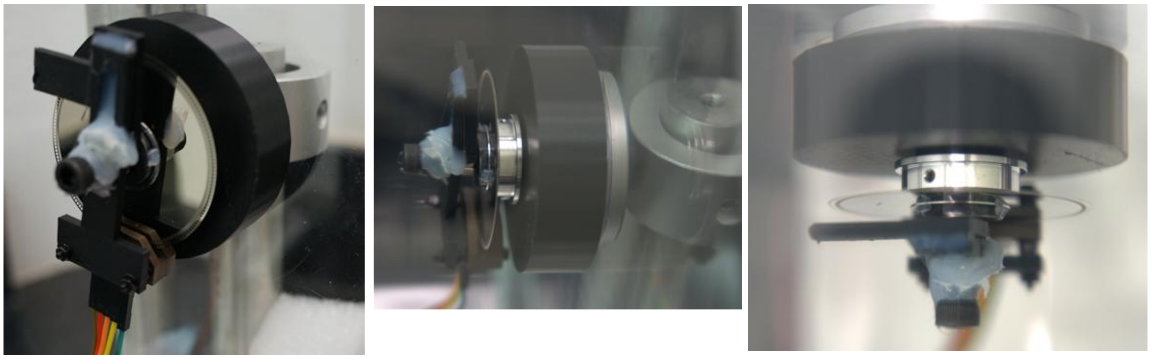

*Figure AG-S2. Rotary encoder coupling assembly used for measuring wheel rotation. Detailed views of the encoder system used to measure wheel rotation. The encoder assembly consists of a rotary encoder wheel, a ball-bearing coupling, and a rotary encoder transceiver module. The encoder wheel is attached to the rotating portion of the bearing so that it rotates synchronously with the transparent running wheel. The encoder transceiver module is mounted to the stationary structure and detects rotational movement of the encoder wheel. A custom coupling interface is used to connect the encoder wheel to the wheel hub. The encoder signal is transmitted through a multi-wire cable connected to the data acquisition system.*

#### Step 4. Head-Fixation Assembly

The head-fixation system is mounted at the top of the 1-inch aluminum rod.

1. Attach a 1-inch rod clamp to the top of the central rod.
2. Mount a ½-inch × ½-inch square aluminum bar using a perpendicular clamp.

This square aluminum bar serves as the headplate holder and contains threaded holes that match the screws used to secure the implanted headplate.

#### Step 5. Animal Casing

A 3D-printed animal casing is mounted to the same square aluminum bar used for the head-fixation assembly.

The casing restricts lateral movement and stabilizes the animal relative to the running wheel and cameras.

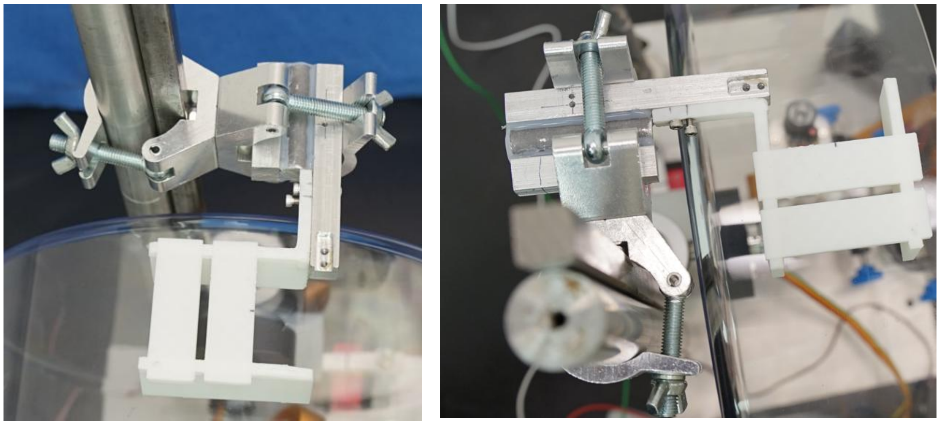

*Figure AG-S3. Head-fixation assembly and animal casing. Detailed view of the head-fixation mechanism mounted on the vertical 1-inch aluminum support rod using adjustable metal rod clamps. A square aluminum crossbar holds the head-fixation bracket that secures the implanted headplate of the animal during experiments. The animal casing, shown here in white, is mounted directly below the fixation bracket and helps stabilize the animal's body relative to the running wheel while allowing free locomotion. In the main manuscript figures the casing is shown in black; the white version shown here represents an alternate printed version of the same component used during prototype assembly.*

#### Step 6. Camera Mounting

Three Raspberry Pi cameras are used:
• Bottom (paw) camera
• Face camera
• Eye-Pupil camera

Side rod camera mounts:
The ½-inch rod on the same side as the head-fixation assembly supports the face and pupil cameras using rod clamps.

The pupil camera is attached using adjustable lever mounts to allow rotational alignment with the eye.

Bottom camera mount:
The ½-inch rod on the opposite side of the base plate supports the bottom-view camera which captures ventral paw movements through the transparent wheel.

Paws and Face cameras use the original Raspberry Pi mounting hardware. The pupil camera uses adjustable lever mounts.

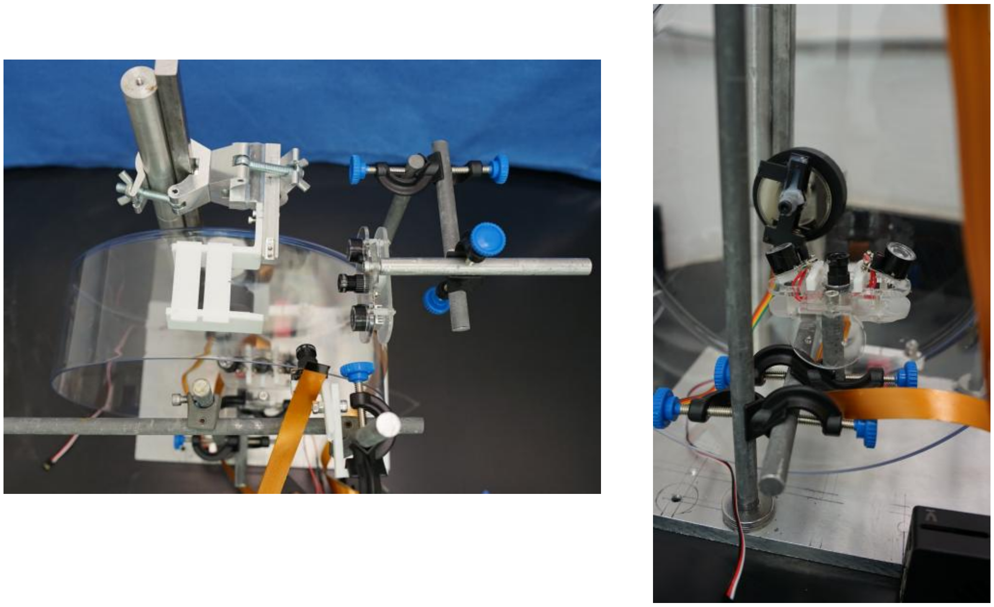

*Figure AG-S4. Mounting configuration for facial and pupil cameras in the AIR-Wheel apparatus. Views of the camera mounting arrangement used for facial and ocular videography during head-fixed locomotion experiments. The cameras are mounted on adjustable aluminum rods using standard optical clamps, allowing precise positioning relative to the animal's head and the transparent running wheel. Left: Top view of the apparatus showing the positioning of the facial and pupil cameras relative to the head-fixation assembly and running wheel. Adjustable rod clamps allow fine alignment of the cameras with respect to the animal's face. Right: Close-up view of the paws camera assembly.*

#### Step 7. Air Delivery System

Air stimulation is delivered using a 3 mL syringe nozzle mounted near the animal's back.

The syringe is mounted using:
• A ½-inch circular aluminum rod
• Standard rod clamps

This rod attaches to the central 1-inch rod. The syringe can be repositioned to direct airflow toward the animal's back. Pneumatic circuit diagram is given below in Figure AG-S5.

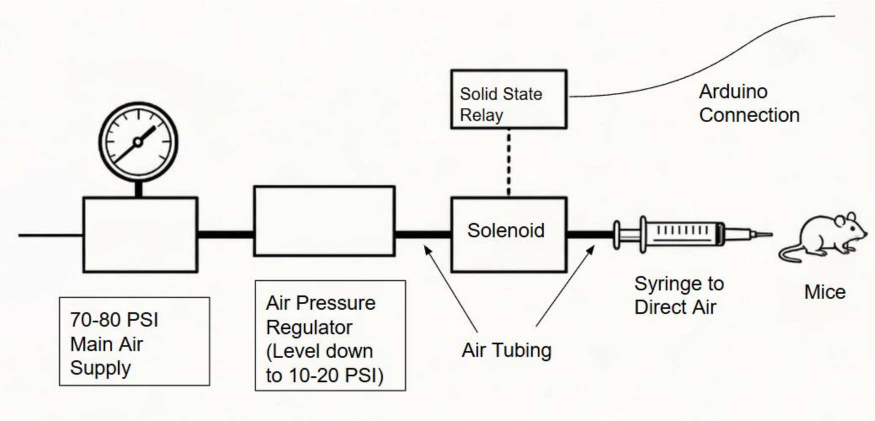


*Figure AG-S5. Pneumatic control system for air delivery in the AIR-Wheel paradigm. Schematic diagram of the pneumatic system used to deliver controlled air stimuli to the animal during AIR-Wheel experiments. Compressed laboratory air (70–80 PSI) is first passed through an air pressure regulator that reduces the pressure to the operating range of approximately 10–20 PSI. The regulated air is routed through air tubing to an electronically controlled solenoid valve. The solenoid is actuated via a solid-state relay connected to an Arduino microcontroller, allowing precise timing of air delivery during experimental trials. Air exiting the solenoid is directed through a syringe nozzle positioned behind the animal to deliver brief air puffs to the animal's back, thereby inducing locomotion on the transparent running wheel.*

#### Step 8. Electrical Connections

Detailed wiring for cameras, encoder signals, and Arduino control is provided in the Electrical Circuit Diagram (Fig. AG-S6). Camera cables connect to Raspberry Pi computers.

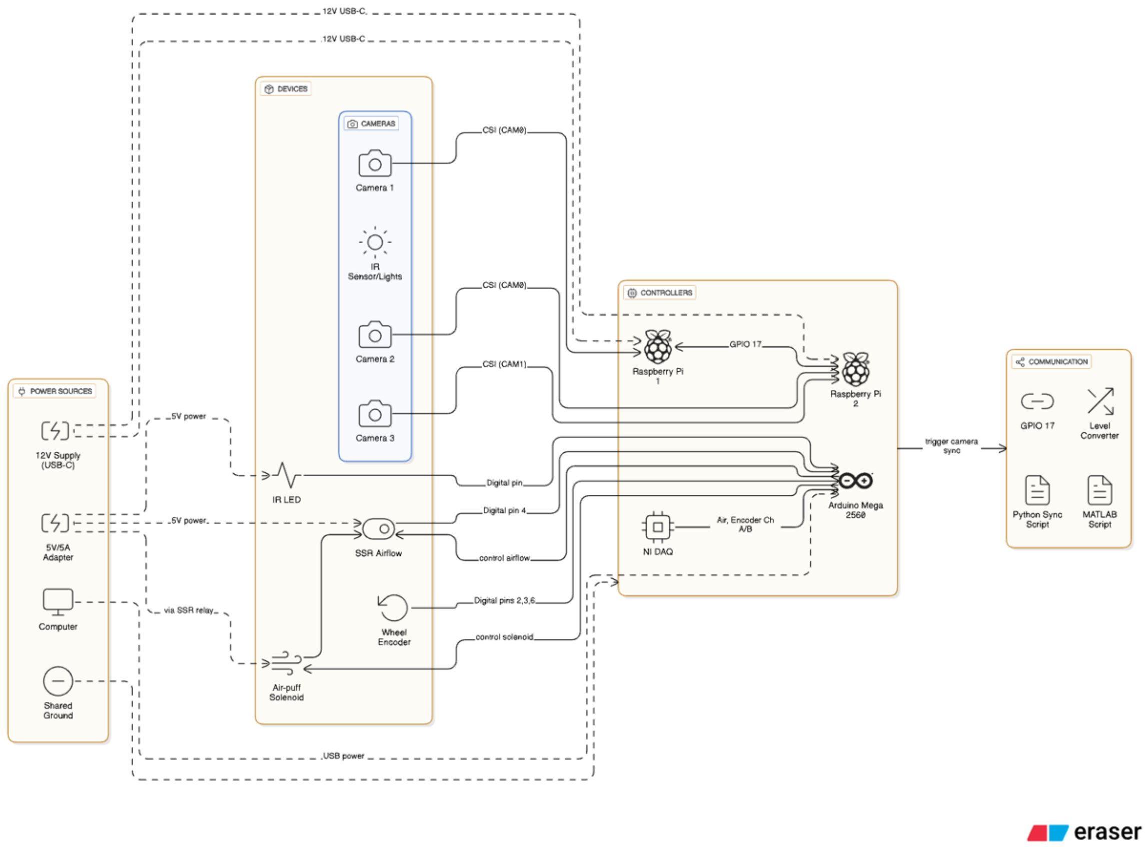


*Figure AG-S6. Electrical and control system architecture of the AIR-Wheel behavioral platform. Schematic diagram illustrating the electrical connections and control architecture used to synchronize behavioral stimulation, wheel movement measurements, and multi-camera video acquisition. The system consists of three major components: power sources, devices, and controllers, interconnected through digital and communication interfaces. Three cameras (Camera 1–3) are connected to two Raspberry Pi controllers through CSI camera interfaces for behavioral video acquisition. Infrared (IR) illumination is provided by an IR LED light source attached to cameras and powered by a regulated 5-V supply to enable imaging under low-light conditions. Wheel rotation is measured using a rotary encoder, which transmits signals through digital input pins to an Arduino Mega 2560 microcontroller. The Arduino also controls the air-puff solenoid valve that delivers airflow stimuli via a solid-state relay (SSR) used to regulate airflow. Encoder signals and airflow timing signals are simultaneously transmitted to a National Instruments (NI) data acquisition (DAQ) system for recording and synchronization with other experimental signals. The Raspberry Pi controllers communicate through GPIO trigger signals (GPIO 17) to synchronize camera acquisition across devices. These trigger signals can be interfaced through a level converter and controlled using either a Python synchronization script or MATLAB-based control scripts.*

#### System Operation and Data Acquisition Workflow

The behavioral system integrates three main components:
(1) Arduino-based stimulus control**,**
(2) Raspberry Pi–based video acquisition, and
(3) MATLAB-based data acquisition using the NI-DAQ card**.**

The Arduino firmware controlling the air-delivery system is provided in the Supplementary Materials and must be uploaded to the Arduino before running the system.

##### Step 1: Initialize the DAQ system

The National Instruments DAQ card is first initialized using the MATLAB initialization script. This script configures the DAQ device, prepares the digital output channel used for triggering, and verifies connectivity with the hardware.

The DAQ card performs two primary functions during experiments:

- Recording analog signals (e.g., air delivery signal)
- Recording quadrature encoder signals (or counter update on DAQ) from the running wheel to compute wheel position and speed

Additional analog or digital channels can also be added to the DAQ configuration if simultaneous signals from electrophysiology or two-photon imaging systems are to be recorded.

##### Step 2: Start Raspberry Pi camera scripts

Each Raspberry Pi runs a Python acquisition script that controls the cameras.

Two scripts are used:

- **Paw camera script** (single camera recording)

  video_recorder.py

- **Facial and pupil camera script** (two cameras with different resolutions)

  video_recorder_2cams_diff_res.py

These scripts should be copied to the Raspberry Pi only once and can then be executed at the start of each experiment.

When launched, the scripts configure the cameras and **wait for a digital trigger on GPIO pin 17**. The cameras remain idle until the trigger signal is received.

##### Step 3: Begin MATLAB recording

The main MATLAB acquisition script is then executed. This script performs the following tasks:

- Starts continuous acquisition from the DAQ card
- Records the air-control signal and wheel encoder position
- Sends a **digital TTL trigger (P0.0)** to the Raspberry Pi systems

When the TTL signal transitions to HIGH, the Raspberry Pi scripts detect the trigger and automatically begin recording video from their cameras.

This ensures that video acquisition and DAQ data acquisition begin simultaneously**.**

##### Step 4: Behavioral recording

During the recording session:

- The DAQ records:
  - Air-delivery signal
  - Wheel encoder counts
- The Raspberry Pi systems record:
  - Bottom-view paw video
  - Facial video
  - Pupil video

All cameras record continuously while the trigger signal remains HIGH.

An LED indicator driven by the same control signal as the air valve is visible in each camera's field of view, allowing precise synchronization of video frames with behavioral and DAQ signals during offline analysis.

##### Step 5: End recording

Recording continues for the duration specified in the MATLAB acquisition script.

When the recording session ends:

1. MATLAB sets the trigger signal LOW.
2. The Raspberry Pi scripts detect the LOW signal.
3. Camera recording automatically stops.
4. Video files are saved on the Raspberry Pi systems.

5. The DAQ acquisition stops and saves the behavioral data to a MATLAB `.mat` file.

##### Step 6: Data output

The system produces the following outputs for each session:

**DAQ recordings**

- Air-delivery signal
- Wheel encoder position and speed
- Optional additional signals (e.g., neural recordings)

**Video recordings**

- Ventral paw camera
- Facial camera
- Pupil camera

All data streams can be aligned offline using the LED synchronization signal embedded in the videos.

#### Data Organization and Loading Workflow

Each experimental session produces a set of raw behavioral and video files that are stored in a structured directory format. This organization ensures that behavioral recordings, video recordings, and analysis outputs remain synchronized and easy to process.

##### Raw Data Structure

For each experiment, a **raw data directory** is created that contains recordings for a specific **animal and session date**.

A typical directory structure is:

```
Raw_Data/
   Animal_ID/
       YYYY_MM_DD/
            recording_YYYYMMDD_HHMMSS.mat
            paws_*.h264
            face_*.h264
            pupil_*.h264
```

Each session therefore contains **four primary files**:

1. **DAQ recording (.mat file)**
   This file contains behavioral signals recorded by the National Instruments DAQ system, including:
   - Air stimulus signal
   - Wheel encoder counts
   - Time vector
   - Sampling rate and channel names
2. **Paw camera video (.h264)**
   Bottom-view video showing paw and body movement.
3. **Face camera video (.h264)**
   Front or side facial video used to analyze facial dynamics.
4. **Pupil camera video (.h264)**
   High-magnification video of the eye region used for pupil analysis.

These files are produced during the recording session by the MATLAB DAQ acquisition script and the Raspberry Pi camera acquisition scripts.

---

#### Data Processing Workflow

After recording, data are processed using MATLAB scripts that load and organize the session data.

##### Step 1: Define project directories

The analysis script first defines the main project directory and the subdirectories used for data storage:

- **Raw Data (RData)** – original recordings
- **Processed Data (PData)** – processed intermediate results
- **Analysis Data (AData)** – final analysis outputs and figures

Example structure:

```
AIR_Wheel_Methods/
    RData/
    PData/
    AData/
```

---

##### Step 2: Load experiment metadata

The script then defines the list of animals and recording dates:

```
animal_list = {'Animal1','Animal2',...}
date_list   = {'YYYY_MM_DD', ...}
```

A helper function (`get_exp_info`) reads this information and generates a structured dataset describing all experiments.

This structure stores:

- Animal identifiers
- Session dates
- File paths to raw recordings
- Paths for processed data

---

#### Step 3: Process behavioral signals

The DAQ recording file is then processed using the function:

```
process_behavior_signals()
```

This step extracts:

- Air stimulus timing
- Encoder counts
- Distance traveled
- Running speed

These variables are stored in the experiment structure for further analysis.

---

#### Step 4: Convert and process videos

The raw `.h264` video files are processed using the function:

```
process_h264()
```

This step:

1. Counts the number of frames in each video.
2. Computes the effective frame rate using the DAQ recording duration.
3. Converts `.h264` recordings to `.mp4` format.
4. Saves video metadata (frame count and frame rate).

The resulting videos are used for computer vision analysis.

##### Step 5: Load LED synchronization signals

LED signals embedded in the videos are extracted using:

```
load_led_signal()
```

These signals identify **air stimulus onset and offset** within the video streams, enabling alignment of behavioral and video data.

##### Step 6: Load pupil region and signals

For experiments that include eye recordings, additional steps load the pupil region of interest (ROI) and extract pupil signals:

```
load_eye_pupil_roi()
load_eye_pupil_signal()
```

These steps allow analysis of pupil dynamics and eye-related behavioral variables.

##### Step 7: Load video files for analysis

The processed video files are then opened in MATLAB using the `VideoReader` function.

Example cameras loaded include:

- Paw camera
- Face camera
- Pupil camera

These videos can then be used for:

- Pose estimation (DeepLabCut)
- Optical flow analysis
- Behavioral visualization

#### Behavioral Signal Processing (MATLAB)

The MATLAB function `process_behavior_signals.m` processes the behavioral data recorded during AIR-wheel experiments. The input to this function is an `animal` structure containing the behavioral recording file generated by the DAQ acquisition script.

The purpose of this script is to convert raw DAQ signals into interpretable behavioral variables such as **air stimulus timing, wheel position, distance traveled, and running speed**.

##### Loading recorded behavioral data

For each experimental session, the script loads the `.mat` file generated by the MATLAB acquisition program. This file contains:

- Recorded analog signals (e.g., air stimulus signal)
- Encoder counts from the running wheel
- Time stamps
- Sampling rate information
- Channel names

These signals are stored in the variable `X`, while the time vector is stored in `t`.

##### Detecting stimulus transitions

The script identifies the timing of behavioral events by detecting **rising and falling edges** in the recorded signals.

For each channel (except the encoder channel), two event types are computed:

- **Rising edges** – transitions from low to high signal values
- **Falling edges** – transitions from high to low signal values

For the air stimulus signal specifically:

- The signal is normalized
- A threshold is applied to generate a **binary air-on / air-off signal**

This binary signal is used to identify stimulus epochs.

##### Encoder signal processing

The running wheel encoder measures wheel rotation. Two possible encoder configurations are supported:

##### Quadrature encoder channels

If channels `ChA` and `ChB` are present, the function reconstructs wheel position using a **quadrature decoding algorithm** implemented in the helper function:

```
processEncodeSignals()
```

This algorithm determines the direction of rotation and increments or decrements the encoder count accordingly.

##### Direct encoder count

If the DAQ system recorded encoder position directly (`EncCount` channel), this value is used directly.

The result is a continuous estimate of **wheel position in encoder counts**.

#### Distance calculation

Wheel position is converted to physical distance using the wheel circumference:

```
distance = encoderCount × (π × wheel_diameter) / countsPerRev
```

In the current implementation the wheel diameter is **32 cm**, producing distance in **centimeters**.

#### Running speed calculation

Running speed is computed as the derivative of distance with respect to time:

```
speed = diff(distance) / diff(time)
```

The first value is set to zero to maintain vector length.

This produces instantaneous running speed in **cm/s**.

#### Speed smoothing

A smoothed speed signal is also computed by applying a moving-average filter over a one-second window:

```
filteredSpeed = moving average of speed
```

This filtered signal is useful for visualizing locomotion dynamics and reducing high-frequency noise.

#### Video Processing and Frame-Rate Correction (MATLAB)

The MATLAB function **`process_h264.m`** converts the raw video recordings generated by the Raspberry Pi cameras into analysis-ready video files and computes accurate timing information for each recording.

During experiments, Raspberry Pi cameras save videos in **H.264 format (`.h264`)**. These files contain raw video streams without a reliable timestamp structure. Because each Raspberry Pi camera runs on its own internal clock, the actual frame rate can differ slightly from the nominal frame rate configured during recording.

This script resolves these issues by computing the **effective frame rate of each video using the DAQ recording duration as the ground-truth time reference**, and then converting the raw video files into properly timed `.mp4` files.

---

##### Processing Workflow

##### 1. Identify recorded video files

For each experimental session, the script checks the `animal` data structure to identify available video recordings from the three cameras:

- **Face camera**
- **Pupil camera**
- **Paw (ventral) camera**

For each camera, the script retrieves the corresponding `.h264` video file recorded by the Raspberry Pi.

---

##### 2. Obtain the true recording duration

The **true recording duration** is obtained from the DAQ behavioral recording.
The DAQ system provides a reliable time reference because it records signals using a fixed sampling rate.

The script extracts the final timestamp from the DAQ time vector:

```
duration = animal(an).b.t(end)
```

This duration represents the actual length of the behavioral recording session.

---

##### 3. Count the number of frames in the video

The script uses the external tool **FFprobe** (part of the FFmpeg software package) to count the number of frames in the raw video file.

This step determines:

- Total number of frames in the `.h264` recording

```
number_of_frames
```

---

##### 4. Compute the effective frame rate

Using the frame count and the true DAQ recording duration, the script computes the **effective frame rate**:

```
effective_fps = number_of_frames / recording_duration
```

Because the Raspberry Pi cameras run on independent clocks, this effective frame rate may differ slightly from the nominal recording rate (e.g., 60 Hz).

Using the DAQ duration ensures that the video timing matches the behavioral recording.

---

##### 5. Convert the raw video file to MP4

The script then converts the `.h264` video file into an `.mp4` container using **FFmpeg**.

During this conversion:

- The computed effective frame rate is applied
- Frame timing information is reconstructed
- The video stream itself is not re-encoded (to preserve quality)

This produces a new video file with correct timing information that can be used for behavioral analysis.

---

#### 6. Save video metadata

For each video file, the script stores the following information:

- Effective frame rate
- Total number of frames

This information is saved in a separate metadata file:

`*_video_specs.mat`

These values are also stored in the `animal` data structure for later use during video analysis and synchronization with behavioral signals.

---

#### Final Output

For each recording session, the script generates:

**Processed video files**

- `face.mp4`
- `pupil.mp4`
- `paws.mp4`

**Video metadata files**

- Effective frame rate
- Frame count

These processed videos are then used for:

- DeepLabCut pose estimation
- Optical-flow motion analysis
- Behavioral synchronization with DAQ signals
