## Supplementary material for "A TRANSPARENT WHEEL-BASED PLATFORM FOR LOCOMOTION-ON-DEMAND AND MULTI-VIEW BODY AND FACIAL KINEMATICS IN HEAD-FIXED MICE": Arduino Code: air_training.ino

```
#include <Servo.h>

#define REWARD_DURATION 100
#define DISTANCE_COVERED 25 // cm
#define START_DISTANCE_COVERED 5// in cm
#define INCREMENT_DISTANCE 1.25 // in cm
#define TOTAL_TRIALS_NB 10000
#define INTERTRIAL_DURATION 15000 // ms
#define SPEED_THRESHOLD 3.5 // in cm/s
#define WHEEL_DIA 32 // in cm
#define PI 3.141592654
#define PPR 4000

Servo myservo;
int servo_pos_NB = 70;
int servo_pos_B = 115;
int servo_pos_R = 70;
int pin_chA = 2; // Interrupt pin for encoder channel A
int pin_chB = 3; // Interrupt pin for encoder channel B
int pin_mode = 6;
int pin_trigger = 5;
int pin_free_rwd = 7;
int pin_free_air = 4 ;

int pin_rwd = 8; // Digital output pin for valve is number 0
int pin_ave = 53; // Digital output pin for aversive stimulus
int pin_stim = 9; // Digital output pin for acoustic stimulus
int pin_light = 20; // Digital output pin for light stimulus
int pin_release_belt = 18;
int pin_photo_sensor = 16;
int pin_trigger_raspi = 34;

long encoder_count = 0;
long last_encoder_count = 0;

int phase = 50;
unsigned int ts = 0;
unsigned int dur = 0;
unsigned int ts1 = 0;
unsigned int dur1 = 0;
unsigned int trials = 0;
unsigned int trialDistance = 0;
float trialDistancecm = START_DISTANCE_COVERED;
unsigned int tsD = 0;
unsigned int durD = 0;
float mouse_speed = 0;
float f_pulses_to_cm = (PI * WHEEL_DIA)/PPR;
float f_cm_to_pulses = PPR/(PI * WHEEL_DIA);

char serialRead = 0;

void setup() {
  // put your setup code here, to run once:
  pinMode(pin_rwd, OUTPUT);
  pinMode(pin_ave, OUTPUT);
  pinMode(pin_stim, OUTPUT);
  pinMode(pin_light, OUTPUT);
  pinMode(pin_trigger_raspi, OUTPUT);


  pinMode(pin_chA, INPUT); // channel A from encoder
  pinMode(pin_chB, INPUT); // channel B from encoder
  pinMode(pin_trigger, INPUT); // trigger input from daq computer
  pinMode(pin_mode, INPUT_PULLUP);
  pinMode(pin_free_rwd, INPUT_PULLUP);
  pinMode(pin_free_air, INPUT_PULLUP);
  pinMode(pin_release_belt, INPUT_PULLUP);
  pinMode(pin_photo_sensor,INPUT);

  digitalWrite(pin_rwd, LOW); // low means off
  digitalWrite(pin_ave, LOW); // low means off
  digitalWrite(pin_stim, LOW); // low means off
  digitalWrite(pin_light, LOW); // low means off
  digitalWrite(pin_trigger_raspi, LOW); // low means off
  // attachInterrupt(digitalPinToInterrupt(pin_chA), encoder_funcAR, RISING); //
  // attachInterrupt(digitalPinToInterrupt(pin_chB), encoder_funcBR, RISING); //
  // attachInterrupt(digitalPinToInterrupt(pin_chA), encoder_funcAF, FALLING); //
  // attachInterrupt(digitalPinToInterrupt(pin_chB), encoder_funcBF, FALLING); //
  attachInterrupt(digitalPinToInterrupt(pin_chA), encoderA, CHANGE);
  attachInterrupt(digitalPinToInterrupt(pin_chB), encoderB, CHANGE);
   myservo.attach(17);
  Serial.begin(115200);
  ts = millis();
}

void loop() {
  // start of the trial
  if (digitalRead(pin_mode) == 0 & digitalRead(pin_trigger) == 1) {
    // Serial.println(encoder_count);
    // No brake loop
    if (phase == 60) {
      trials++;
      if (trials <= TOTAL_TRIALS_NB){
        digitalWrite(pin_trigger_raspi, HIGH); // low means off
         digitalWrite(pin_stim, HIGH); // low means off
         //tone(pin_stim, 6000);  // Generate 1 kHz tone
        phase = 100;
          }
      else {
        trials = 0;
        phase = 1000;
        ts = millis();
      }
    }
    if (phase == 100) {
      myservo.write(servo_pos_NB);
      delay(100);
      phase = 200;
    }
    if (phase == 200) {
      //Serial.println(pin_mode);
      Serial.print("Trial Number: ");
      Serial.print(trials);
      Serial.print("         ");
      Serial.print("Distance: ");
      Serial.print(trialDistancecm);
      Serial.print(" cm             ");
      // Serial.println(trialDistance);
      digitalWrite(pin_ave, HIGH);
      encoder_count = 0;
      tsD = millis();
      phase = 320;
    }
    if (phase == 320) {
      if (encoder_count < 0)
        encoder_count = 0;
        if (encoder_count > trialDistance){
        digitalWrite(pin_ave, LOW);
        phase = 600;
        ts = millis();
        durD = millis()-tsD;
        mouse_speed = trialDistancecm*1000/durD; // speed in cm/sec
        Serial.print("Speed: ");
        Serial.print(mouse_speed);
        Serial.println(" cm/s");
        Serial.println(" ");
        }
    }
  if (phase == 400) {
      dur = millis() - ts;
      if (dur > 10) {
        digitalWrite(pin_rwd, HIGH);
        phase = 500;
        ts1 = millis();
      }
    }
    if (phase == 500) {
      dur = millis() - ts1;
      if (dur > REWARD_DURATION) {
        digitalWrite(pin_rwd, LOW);
        phase = 600;
      }
    }
    if (phase == 600) {
      dur = millis() - ts;
      if (dur > INTERTRIAL_DURATION) {
        phase = 60;
        if (mouse_speed>=SPEED_THRESHOLD)
        trialDistancecm = trialDistancecm + INCREMENT_DISTANCE;
//        if (mouse_speed<SPEED_THRESHOLD)
//          trialDistance = trialDistance - INCREMENT_DISTANCE;
        if (trialDistancecm>DISTANCE_COVERED)
          trialDistancecm = DISTANCE_COVERED;
        if (trialDistancecm<START_DISTANCE_COVERED)
          trialDistancecm = START_DISTANCE_COVERED;
        trialDistance = trialDistancecm * f_cm_to_pulses;
      }
    }
  }
  if (digitalRead(pin_mode) == 1 || digitalRead(pin_trigger) == 0) {
    trials = 0;
    encoder_count = 0;
    digitalWrite(pin_trigger_raspi, LOW); // low means off
    digitalWrite(pin_stim, LOW); // low means off
     //noTone(pin_stim);      // Stop sound
    trialDistance = trialDistancecm * f_cm_to_pulses;
    if (digitalRead(pin_free_rwd) == 0)
      myservo.write(servo_pos_B);
    else
      myservo.write(servo_pos_NB);
    digitalWrite(pin_rwd, LOW); // low means off
    digitalWrite(pin_ave, LOW); // low means off
    digitalWrite(pin_stim, LOW); // low means off
    digitalWrite(pin_light, LOW);
    phase = 60;
    
//    if (digitalRead(pin_release_belt) == 0)
//      myservo.write(servo_pos_R);
//    else
//      myservo.write(servo_pos_B);
//    
    delay(15);
    ts = millis();
//    Serial.println(4);
//    if (digitalRead(pin_free_rwd) == 0) {
//      digitalWrite(pin_rwd, HIGH); //
//      delay(REWARD_DURATION);
//      digitalWrite(pin_rwd, LOW);
//      delay(500);
//    }
    if (digitalRead(pin_free_air) == 0) {
      digitalWrite(pin_ave, HIGH); //
      delay(3000);
    }
    if (digitalRead(pin_photo_sensor) == 0) {
      if (encoder_count > 3000){
      digitalWrite(pin_rwd, HIGH); //
      delay(REWARD_DURATION);
      digitalWrite(pin_rwd, LOW);
      delay(500);
      encoder_count = 0;
      }
    }
  }
}

void encoderA() {
  // Channel A changed
  if (digitalRead(pin_chA) == digitalRead(pin_chB))
    encoder_count++;
  else
    encoder_count--;
}

void encoderB() {
  // Channel B changed
  if (digitalRead(pin_chA) != digitalRead(pin_chB))
    encoder_count++;
  else
    encoder_count--;
}
```
