## Supplementary material for "A TRANSPARENT WHEEL-BASED PLATFORM FOR LOCOMOTION-ON-DEMAND AND MULTI-VIEW BODY AND FACIAL KINEMATICS IN HEAD-FIXED MICE": Arduino Code: habituation.ino

```
#define ON_SECONDS   15
#define OFF_SECONDS  15

const int pin_ave  = 53;
const int pin_mode = 6;   // INPUT_PULLUP, ON -> LOW

enum State { AIR_ON, AIR_OFF };
State state = AIR_OFF;

unsigned long lastToggleMs = 0;
bool wasModeOn = false;

void setAirAndLed(bool on) {
  digitalWrite(pin_ave, on ? HIGH : LOW);
}

void resetPulserToOff() {
  state = AIR_OFF;
  setAirAndLed(false);
  lastToggleMs = millis();
}

void setup() {
  pinMode(pin_ave, OUTPUT);
  digitalWrite(pin_ave, LOW);

  pinMode(pin_mode, INPUT_PULLUP);

  Serial.begin(115200);
  resetPulserToOff();
}

void loop() {
  bool modeOn = (digitalRead(pin_mode) == LOW);

  // Mode OFF: force OFF once, then idle
  if (!modeOn) {
    if (wasModeOn) {
      resetPulserToOff();
      Serial.println("Mode OFF -> forced OFF");
    }
    wasModeOn = false;
    delay(10);
    return;
  }

  // Mode just turned ON: reset timing and start in OFF (as you want)
  if (!wasModeOn) {
    resetPulserToOff(); // starts OFF, waits OFF_SECONDS
    Serial.println("Mode ON -> pulser running");
    wasModeOn = true;
  }

  unsigned long now = millis();
  unsigned long intervalMs =
      (state == AIR_ON) ? (unsigned long)ON_SECONDS * 1000UL
                        : (unsigned long)OFF_SECONDS * 1000UL;

  if (now - lastToggleMs >= intervalMs) {
    state = (state == AIR_ON) ? AIR_OFF : AIR_ON;
    setAirAndLed(state == AIR_ON);
    lastToggleMs = now;

    if (state == AIR_ON) Serial.println("AIR+LED: ON");
    else                 Serial.println("AIR+LED: OFF");
  }
}
```
