## Supplementary material for "A TRANSPARENT WHEEL-BASED PLATFORM FOR LOCOMOTION-ON-DEMAND AND MULTI-VIEW BODY AND FACIAL KINEMATICS IN HEAD-FIXED MICE": Matlab Code: air_training_01.m

```
%% === TTL START on P0.0 (separate object) ===
dev = "Dev1";

recordMinutes = 22;
recordSeconds = recordMinutes * 60;

dDO = daq("ni");                                   % Digital OUTPUT only

addoutput(dDO, dev, "port0/line0", "Digital");     % P0.0 digital out

% Create a new log file
ts = datetime('now','Format','yyyyMMdd_HHmmss');
fname = sprintf('recording_%s.mat', string(ts));

username = getenv("USERNAME");   % e.g., "paranjap"
path = fullfile("C:","Users",username,"data");
if ~exist(path,"dir")
    mkdir(path);
end
fname = fullfile(path,fname);

% === CONTINUOUS ACQUISITION: AIR (AI0) + ENCODER COUNTS (CTR1) ===
fs  = 5000;                      % sample rate (Hz)
d = daq("ni");                   % ONE DAQ object (AI + Counter)

% Air signal only
addinput(d, dev, "ai0", "Voltage");   % Air
d.Rate = fs;

% Encoder counter position on CTR1
enc = addinput(d, dev, "ctr1", "Position");
enc.EncoderType  = "X4";
enc.InitialCount = 0;

% OPTIONAL: enable Z reset if you want (only if Z wired)
enc.ZResetEnable    = true;
enc.ZResetValue     = 0;
enc.ZResetCondition = "AHigh";   % or BHigh/BothHigh/BothLow

disp("Encoder wiring terminals (CTR1):");
disp("A -> " + enc.TerminalA);
disp("B -> " + enc.TerminalB);
disp("Z -> " + enc.TerminalZ + " (optional)");

% === Display-only wrap settings ===
countsPerRev = 4000;   % <--- your effective counts/rev with X4

% Prepare MAT-file logger
m = matfile(fname, 'Writable', true);
m.fs = fs;
m.device = dev;
m.startTime = datetime('now');
m.channelNames = {'Air','EncCount'};   % EncCount saved as TRUE signed counts
m.countsPerRev = countsPerRev;
nTotal = 0;

% === Rolling Buffer for Plot (10 s window) ===
bufferSeconds = 10;
N = fs * bufferSeconds;
buf = zeros(N,2,'single');        % Air + EncCountDisplayed
tPlot = (0:N-1)' / fs;

figure(101); clf;
ax = axes('Parent', gcf);
plt = plot(ax, tPlot, buf);
legend(ax, {'Air','EncCount(mod)'}, 'Location','northwest');
xlabel(ax,'Time (s)');
ylabel(ax,'Value');
title(ax, sprintf('Logging to %s', fname), 'Interpreter','none');
grid(ax,'on');
ylim([-5000 5000])

% === Send TTL START (P0.0 HIGH) ===
write(dDO, true);               % HIGH start trigger
pause(0.01);                    % small settling delay

% === Start continuous acquisition ===
start(d, "continuous");
disp(['Recording (AI0 + CTR1 Position) ... saving to ', fname]);
disp('Close the figure or press Ctrl+C to stop.');

t0 = [];

try
    tStartWall = tic;
    while ishandle(ax) && toc(tStartWall) < recordSeconds

        TT = read(d, seconds(0.1));    % ~0.1 s chunk
        if isempty(TT)
            drawnow;
            continue;
        end

        % Establish t0 at first chunk
        if isempty(t0)
            t0 = TT.Time(1);
        end

        % Convert timetable times to relative seconds
        relT = seconds(TT.Time - t0);

        % Column 1: Air (ai0)
        air = single(TT{:,1});

        % Column 2: Encoder position (ctr1) - convert to signed int32
        posRaw = TT{:,2};
        if isa(posRaw,'uint32')
            posSigned = double(typecast(posRaw, 'int32'));
        else
            posSigned = double(posRaw);
            posSigned(posSigned > 2^31) = posSigned(posSigned > 2^31) - 2^32;
        end
        posSigned = single(posSigned);

        % --- DISPLAY ONLY: wrap to 0..countsPerRev-1 ---
        posDisp = mod(posSigned, countsPerRev);
        posDisp(posDisp < 0) = posDisp(posDisp < 0) + countsPerRev;  % handle reverse

        % ---- Save TRUE signed counts (posSigned), not the wrapped value ----
        chunkSave = [air, posSigned];
        k = size(chunkSave,1);

        m.X(nTotal+1:nTotal+k,1:2) = chunkSave;
        m.t(nTotal+1:nTotal+k,1)   = single(relT);
        nTotal = nTotal + k;

        % ---- Update rolling plot: Air + wrapped encoder for display ----
        airD = air*4000/5;
        chunkPlot = [airD, posDisp];

        if k >= N
            buf = chunkPlot(end-N+1:end, :);
        else
            buf = [buf(k+1:end,:); chunkPlot];
        end
        set(plt(1),'YData',buf(:,1));
        set(plt(2),'YData',buf(:,2));
        drawnow limitrate nocallbacks;
    end

catch ME
    warning("Loop aborted: %s", ME.message);
end

% === Cleanup ===
write(dDO, false);        % pull P0.0 LOW at end
disp('Sent low signal to Raspberry Pi computers')

stop(d);
clear d dDO;
disp(['Stopped. File saved: ', fname]);
```
