## Supplementary material for "A TRANSPARENT WHEEL-BASED PLATFORM FOR LOCOMOTION-ON-DEMAND AND MULTI-VIEW BODY AND FACIAL KINEMATICS IN HEAD-FIXED MICE": Matlab Code: initialize.m

```
%% === TTL START on P0.0 (separate object) ===
dev = "Dev1";

dDO = daq("ni");            
% Digital OUTPUT only
addoutput(dDO, dev, "port0/line0", "Digital");     % P0.0 digital out

% === Send TTL START (P0.0 HIGH) ===
write(dDO, false);               % HIGH start trigger
pause(0.01); 
```
