## Supplementary material for "A TRANSPARENT WHEEL-BASED PLATFORM FOR LOCOMOTION-ON-DEMAND AND MULTI-VIEW BODY AND FACIAL KINEMATICS IN HEAD-FIXED MICE": Raspberry Pi Code: Exp_1_video_recorder.py

```
#!/usr/bin/env python3
import lgpio
import subprocess
import time
from datetime import datetime
import os
import signal

TRIGGER_PIN = 17
OUTPUT_DIR = "/home/neuromomentum/recordings"

# Fixed camera parameters
WIDTH = 1280
HEIGHT = 720
FRAMERATE = 60
BITRATE = 6000000
CODEC = "h264"

# Ensure output directory exists
os.makedirs(OUTPUT_DIR, exist_ok=True)

# ----- GPIO Setup -----
chip = lgpio.gpiochip_open(0)
lgpio.gpio_claim_input(chip, TRIGGER_PIN)

print(f"Slave ready - waiting for HIGH on GPIO {TRIGGER_PIN}")

try:
    # Wait until line goes HIGH
    while lgpio.gpio_read(chip, TRIGGER_PIN) == 0:
        time.sleep(0.001)

    # Timestamp for file naming
    timestamp = datetime.now().strftime("%Y%m%d_%H%M%S")
    # outfile = os.path.join(OUTPUT_DIR, f"video_{timestamp}.h264")
    outfile = os.path.join(OUTPUT_DIR, f"video_{WIDTH}x{HEIGHT}_{FRAMERATE}_{timestamp}.h264")

    print(f"Trigger HIGH detected - starting recording ? {outfile}")

    # Start indefinite recording (until SIGINT)
    cmd = [
        "rpicam-vid",
        "--width", str(WIDTH),
        "--height", str(HEIGHT),
        "--framerate", str(FRAMERATE),
 #       "--bitrate", str(BITRATE),
        "--codec", CODEC,
        "-t", "0",              # run until stopped
        "-o", outfile
    ]

    proc = subprocess.Popen(cmd)
    print("Recording... waiting for trigger LOW to stop.")

    # Keep recording while HIGH
    while lgpio.gpio_read(chip, TRIGGER_PIN) == 1:
        if proc.poll() is not None:  # If process died
            print("Camera recording stopped unexpectedly.")
            break
        time.sleep(0.001)

    print("Trigger LOW detected - stopping recording")
    proc.send_signal(signal.SIGINT)
    proc.wait()

    print(f"Recording saved ? {outfile}")

finally:
    lgpio.gpiochip_close(chip)
```
