## Supplementary material for "A TRANSPARENT WHEEL-BASED PLATFORM FOR LOCOMOTION-ON-DEMAND AND MULTI-VIEW BODY AND FACIAL KINEMATICS IN HEAD-FIXED MICE": Raspberry Pi Code: Exp_1_video_recorder_2cams_diff_res.py

```
#!/usr/bin/env python3
import lgpio
import subprocess
import time
from datetime import datetime
import os
import signal

TRIGGER_PIN = 17
OUTPUT_DIR = "/home/neuromomentum/recordings"

# Camera-specific configuration
CAMERA_CONFIG = {
    0: {"name": 'face', "width": 960, "height": 720, "framerate": 60, "bitrate": 5000000, "rot": 180},  # behavioral camera
    1: {"name": 'pupi', "width": 320, "height": 240, "framerate": 60, "bitrate": 3000000, "rot": 0},  # pupil camera
}

CODEC = "h264"   # output container will be MP4

# Ensure output directory exists
os.makedirs(OUTPUT_DIR, exist_ok=True)

# ----- GPIO Setup -----
chip = lgpio.gpiochip_open(0)
lgpio.gpio_claim_input(chip, TRIGGER_PIN)

print(f"Slave ready - waiting for HIGH on GPIO {TRIGGER_PIN}")

try:
    # Wait for rising edge (HIGH)
    while lgpio.gpio_read(chip, TRIGGER_PIN) == 0:
        time.sleep(0.001)

    # Shared timestamp for all camera files
    timestamp = datetime.now().strftime("%Y%m%d_%H%M%S")

    procs = []
    outfiles = []

    # Start recording for each camera
    for cam_idx, cfg in CAMERA_CONFIG.items():
        outfile = os.path.join(
            OUTPUT_DIR,
            f"{cfg['name']}_{cfg['width']}x{cfg['height']}_{cfg['framerate']}_{timestamp}.h264"
        )
        outfiles.append(outfile)

        print(f"Starting camera {cam_idx} ? {outfile}")

        cmd = [
            "rpicam-vid",
            "--camera", str(cam_idx),
            "--width", str(cfg["width"]),
            "--height", str(cfg["height"]),
            "--framerate", str(cfg["framerate"]),
            "--bitrate", str(cfg["bitrate"]),
            "--rot", str(cfg["rot"]),
            "--codec", CODEC,
            "-t", "0",          # run until SIGINT
            "-o", outfile,
            #"--nopreview"       # maximize performance
        ]

        proc = subprocess.Popen(cmd)
        procs.append(proc)

    print("Recording on all cameras... waiting for LOW to stop.")

    # Keep recording while HIGH
    while lgpio.gpio_read(chip, TRIGGER_PIN) == 1:
        # Check if any camera process died unexpectedly
        for cam_idx, proc in zip(CAMERA_CONFIG.keys(), procs):
            if proc.poll() is not None:
                print(f"WARNING: Camera {cam_idx} process exited unexpectedly!")
        time.sleep(0.001)

    print("Trigger LOW detected -stopping all recordings")

    # Stop all camera processes gracefully
    for proc in procs:
        if proc.poll() is None:
            proc.send_signal(signal.SIGINT)

    # Wait for clean shutdown
    for cam_idx, proc in zip(CAMERA_CONFIG.keys(), procs):
        try:
            proc.wait(timeout=10)
        except subprocess.TimeoutExpired:
            print(f"Camera {cam_idx} did not exit in time - killing it.")
            proc.kill()

    # File outputs
    for outfile in outfiles:
        print(f"Saved recording ? {outfile}")

finally:
    lgpio.gpiochip_close(chip)
    print("GPIO chip closed. Script complete.")
```
