## Supplementary material for "A TRANSPARENT WHEEL-BASED PLATFORM FOR LOCOMOTION-ON-DEMAND AND MULTI-VIEW BODY AND FACIAL KINEMATICS IN HEAD-FIXED MICE": Raspberry Pi Code: video_recorder_2cams_diff_res_v2.py

```
#!/usr/bin/env python3
import lgpio
import subprocess
import time
from datetime import datetime
import os
import signal
import argparse
import sys

TRIGGER_PIN = 17

# Default output dir (change with --outdir)
DEFAULT_OUTPUT_DIR = "/home/neuromomentum/recordings"

# Camera-specific configuration (you can tweak per cam)
# IMPORTANT:
#  - At 60 fps, shutter MUST be <= 16666 us (frame period 16.67 ms)
#  - At 45 fps, shutter MUST be <= 22222 us (frame period 22.22 ms)
CAMERA_CONFIG = {
    0: {
        "name": "face",
        "width": 1440,
        "height": 1080,
        "framerate": 60,
        "bitrate": 8000000,
        "rot": 180,
        "shutter_us": 10000,   # <-- key fix
        "gain": 4.0,
    },
    1: {
        "name": "pupi",
        "width": 320,
        "height": 240,
        "framerate": 60,
        "bitrate": 5000000,
        "rot": 0,
        "shutter_us": 10000,   # <-- key fix
        "gain": 4.0,
    },
}

CODEC = "h264"  # rpicam-vid will write raw .h264 by default when -o ends with .h264


def parse_args():
    ap = argparse.ArgumentParser(
        description="GPIO-triggered dual-camera rpicam recorder (v2: locks shutter/gain/AWB to prevent frame drops)."
    )
    ap.add_argument("--outdir", default=DEFAULT_OUTPUT_DIR, help="Output directory.")
    ap.add_argument("--framerate", type=int, default=None, help="Override framerate for ALL cameras (e.g., 45 or 60).")
    ap.add_argument("--shutter-us", type=int, default=None, help="Override shutter (microseconds) for ALL cameras.")
    ap.add_argument("--gain", type=float, default=None, help="Override gain for ALL cameras.")
    ap.add_argument("--awb-off", action="store_true", help="Disable AWB (recommended for IR stability).")
    ap.add_argument("--awbgains", default="1.6,1.6", help="AWB gains if --awb-off (format: R,B).")
    ap.add_argument("--nopreview", action="store_true", help="Enable --nopreview for performance.")
    ap.add_argument("--codec", default=CODEC, help="Codec for rpicam-vid (default: h264).")
    return ap.parse_args()


def sanity_check(cfg):
    fps = int(cfg["framerate"])
    shutter = int(cfg["shutter_us"])
    max_shutter = int(1_000_000 / fps)  # microseconds
    if shutter > max_shutter:
        print(
            f"WARNING: {cfg['name']} shutter_us={shutter} is > max allowed for {fps} fps ({max_shutter} us). "
            f"This can force fps to drop / frames to be skipped. Lower shutter-us.",
            file=sys.stderr
        )


def main():
    args = parse_args()

    outdir = args.outdir
    os.makedirs(outdir, exist_ok=True)

    # Apply global overrides
    for cam_idx, cfg in CAMERA_CONFIG.items():
        if args.framerate is not None:
            cfg["framerate"] = int(args.framerate)
        if args.shutter_us is not None:
            cfg["shutter_us"] = int(args.shutter_us)
        if args.gain is not None:
            cfg["gain"] = float(args.gain)
        sanity_check(cfg)

    # ----- GPIO Setup -----
    chip = lgpio.gpiochip_open(0)
    lgpio.gpio_claim_input(chip, TRIGGER_PIN)

    print(f"Slave ready - waiting for HIGH on GPIO {TRIGGER_PIN}")

    try:
        # Wait for rising edge (HIGH)
        while lgpio.gpio_read(chip, TRIGGER_PIN) == 0:
            time.sleep(0.001)

        timestamp = datetime.now().strftime("%Y%m%d_%H%M%S")
        procs = []
        outfiles = []

        # Start recording for each camera
        for cam_idx, cfg in CAMERA_CONFIG.items():
            outfile = os.path.join(
                outdir,
                f"{cfg['name']}_{cfg['width']}x{cfg['height']}_{cfg['framerate']}_{timestamp}.h264"
            )
            outfiles.append(outfile)

            print(f"Starting camera {cam_idx} -> {outfile}")

            cmd = [
                "rpicam-vid",
                "--camera", str(cam_idx),
                "--width", str(cfg["width"]),
                "--height", str(cfg["height"]),
                "--framerate", str(cfg["framerate"]),
                "--bitrate", str(cfg["bitrate"]),
                "--rot", str(cfg["rot"]),
                "--codec", str(args.codec),

                # Key stability/performance flags:
                "--shutter", str(cfg["shutter_us"]),
                "--gain", str(cfg["gain"]),
            ]

            if args.awb_off:
                cmd += ["--awb", "off", "--awbgains", str(args.awbgains)]

            if args.nopreview:
                cmd += ["--nopreview"]

            cmd += [
                "-t", "0",  # run until SIGINT
                "-o", outfile,
            ]

            proc = subprocess.Popen(cmd)
            procs.append(proc)

        print("Recording on all cameras... waiting for LOW to stop.")

        # Keep recording while HIGH
        while lgpio.gpio_read(chip, TRIGGER_PIN) == 1:
            for cam_idx, proc in zip(CAMERA_CONFIG.keys(), procs):
                if proc.poll() is not None:
                    print(f"WARNING: Camera {cam_idx} process exited unexpectedly!")
            time.sleep(0.001)

        print("Trigger LOW detected - stopping all recordings")

        # Stop all camera processes gracefully
        for proc in procs:
            if proc.poll() is None:
                proc.send_signal(signal.SIGINT)

        # Wait for clean shutdown
        for cam_idx, proc in zip(CAMERA_CONFIG.keys(), procs):
            try:
                proc.wait(timeout=10)
            except subprocess.TimeoutExpired:
                print(f"Camera {cam_idx} did not exit in time - killing it.")
                proc.kill()

        for outfile in outfiles:
            print(f"Saved recording -> {outfile}")

    finally:
        lgpio.gpiochip_close(chip)
        print("GPIO chip closed. Script complete.")


if __name__ == "__main__":
    main()
```
