## Supplementary material for "A TRANSPARENT WHEEL-BASED PLATFORM FOR LOCOMOTION-ON-DEMAND AND MULTI-VIEW BODY AND FACIAL KINEMATICS IN HEAD-FIXED MICE": Raspberry Pi Code: video_recorder_paws_v2.py

```
#!/usr/bin/env python3
import lgpio
import subprocess
import time
from datetime import datetime
import os
import signal
import argparse
import sys

TRIGGER_PIN = 17
DEFAULT_OUTPUT_DIR = "/home/neuromomentum/recordings"

# Paws camera config (edit if needed)
CAMERA_CONFIG = {
    0: {
        "name": "paws",          # <-- changed from video to paws
        "width": 1440,
        "height": 1080,
        "framerate": 45,         # safer default
        "bitrate": 6000000,
        "rot": 0,
        "shutter_us": 12000,     # must be <= 22222 for 45 fps
        "gain": 4.0,
    }
}

CODEC = "h264"


def parse_args():
    ap = argparse.ArgumentParser(description="GPIO-triggered paws recorder (v2).")
    ap.add_argument("--outdir", default=DEFAULT_OUTPUT_DIR)
    ap.add_argument("--framerate", type=int, default=None)
    ap.add_argument("--shutter-us", type=int, default=None)
    ap.add_argument("--gain", type=float, default=None)
    ap.add_argument("--awb-off", action="store_true")
    ap.add_argument("--awbgains", default="1.6,1.6")
    ap.add_argument("--nopreview", action="store_true")
    return ap.parse_args()


def sanity_check(cfg):
    fps = int(cfg["framerate"])
    shutter = int(cfg["shutter_us"])
    max_shutter = int(1_000_000 / fps)
    if shutter > max_shutter:
        print(
            f"WARNING: shutter_us={shutter} too high for {fps} fps (max {max_shutter})",
            file=sys.stderr
        )


def main():
    args = parse_args()

    outdir = args.outdir
    os.makedirs(outdir, exist_ok=True)

    # Apply overrides
    for cam_idx, cfg in CAMERA_CONFIG.items():
        if args.framerate is not None:
            cfg["framerate"] = args.framerate
        if args.shutter_us is not None:
            cfg["shutter_us"] = args.shutter_us
        if args.gain is not None:
            cfg["gain"] = args.gain
        sanity_check(cfg)

    chip = lgpio.gpiochip_open(0)
    lgpio.gpio_claim_input(chip, TRIGGER_PIN)

    print(f"Paws recorder ready - waiting for HIGH on GPIO {TRIGGER_PIN}")

    try:
        while lgpio.gpio_read(chip, TRIGGER_PIN) == 0:
            time.sleep(0.001)

        timestamp = datetime.now().strftime("%Y%m%d_%H%M%S")
        procs = []
        outfiles = []

        for cam_idx, cfg in CAMERA_CONFIG.items():

            outfile = os.path.join(
                outdir,
                f"paws_{cfg['width']}x{cfg['height']}_{cfg['framerate']}_{timestamp}.h264"
            )

            outfiles.append(outfile)

            print(f"Starting paws camera {cam_idx} -> {outfile}")

            cmd = [
                "rpicam-vid",
                "--camera", str(cam_idx),
                "--width", str(cfg["width"]),
                "--height", str(cfg["height"]),
                "--framerate", str(cfg["framerate"]),
                "--bitrate", str(cfg["bitrate"]),
                "--rot", str(cfg["rot"]),
                "--codec", CODEC,
                "--shutter", str(cfg["shutter_us"]),
                "--gain", str(cfg["gain"]),
            ]

            if args.awb_off:
                cmd += ["--awb", "off", "--awbgains", args.awbgains]

            if args.nopreview:
                cmd += ["--nopreview"]

            cmd += [
                "-t", "0",
                "-o", outfile,
            ]

            proc = subprocess.Popen(cmd)
            procs.append(proc)

        print("Recording paws... waiting for LOW to stop.")

        while lgpio.gpio_read(chip, TRIGGER_PIN) == 1:
            time.sleep(0.001)

        print("Stopping paws recording")

        for proc in procs:
            if proc.poll() is None:
                proc.send_signal(signal.SIGINT)

        for proc in procs:
            proc.wait(timeout=10)

        for outfile in outfiles:
            print(f"Saved -> {outfile}")

    finally:
        lgpio.gpiochip_close(chip)
        print("Done.")


if __name__ == "__main__":
    main()
```
