## Supplementary material for "A TRANSPARENT WHEEL-BASED PLATFORM FOR LOCOMOTION-ON-DEMAND AND MULTI-VIEW BODY AND FACIAL KINEMATICS IN HEAD-FIXED MICE": Raspberry Pi Code: video_recorder_2cams.py

```
#!/usr/bin/env python3
import lgpio
import subprocess
import time
from datetime import datetime
import os
import signal

TRIGGER_PIN = 17
OUTPUT_DIR = "/home/neuromomentum/recordings"

# Fixed camera parameters
WIDTH = 1920
HEIGHT = 1080
FRAMERATE = 60
BITRATE = 10000000
CODEC = "h264"   # the codec; MP4 is chosen by file extension

CAMERA_CONFIG = {
    0: {"width": 1280, "height": 960, "bitrate": 8000000},   # behavioral
    1: {"width": 1024, "height": 768, "bitrate": 5000000},   # pupil
}


# Cameras to record from on this Pi
CAMERA_INDICES = [0, 1]   # two cameras: 0 and 1

# Ensure output directory exists
os.makedirs(OUTPUT_DIR, exist_ok=True)

# ----- GPIO Setup -----
chip = lgpio.gpiochip_open(0)
lgpio.gpio_claim_input(chip, TRIGGER_PIN)

print(f"Slave ready - waiting for HIGH on GPIO {TRIGGER_PIN}")

try:
    # Wait until line goes HIGH
    while lgpio.gpio_read(chip, TRIGGER_PIN) == 0:
        time.sleep(0.001)

    # Shared timestamp for all cameras on this Pi
    timestamp = datetime.now().strftime("%Y%m%d_%H%M%S")

    procs = []
    outfiles = []

    # Start recording on all cameras
    for cam_idx in CAMERA_INDICES:
        outfile = os.path.join(
            OUTPUT_DIR,
            f"slave_cam{cam_idx}_{timestamp}.mp4"  # MP4 container
        )
        outfiles.append(outfile)

        print(f"Starting recording on camera {cam_idx} ? {outfile}")

        cmd = [
            "rpicam-vid",
            "--camera", str(cam_idx),
            "--width", str(WIDTH),
            "--height", str(HEIGHT),
            "--framerate", str(FRAMERATE),
            "--bitrate", str(BITRATE),
            "--codec", CODEC,
            "-t", "0",              # run until stopped
            "-o", outfile
        ]

        proc = subprocess.Popen(cmd)
        procs.append(proc)

    print("Recording on all cameras... waiting for trigger LOW to stop.")

    # Keep recording while trigger is HIGH
    while lgpio.gpio_read(chip, TRIGGER_PIN) == 1:
        # If any process dies unexpectedly, break
        for cam_idx, proc in zip(CAMERA_INDICES, procs):
            if proc.poll() is not None:
                print(f"Camera process {cam_idx} exited unexpectedly.")
        time.sleep(0.001)

    print("Trigger LOW detected - stopping all recordings")

    # Politely stop all camera processes
    for proc in procs:
        if proc.poll() is None:
            proc.send_signal(signal.SIGINT)

    # Wait for all to exit
    for cam_idx, proc in zip(CAMERA_INDICES, procs):
        try:
            proc.wait(timeout=10)
        except subprocess.TimeoutExpired:
            print(f"Camera process {cam_idx} did not exit in time, killing it.")
            proc.kill()

    for outfile in outfiles:
        print(f"Recording saved ? {outfile}")

finally:
    lgpio.gpiochip_close(chip)
```
