## Supplementary figures and images for "A TRANSPARENT WHEEL-BASED PLATFORM FOR LOCOMOTION-ON-DEMAND AND MULTI-VIEW BODY AND FACIAL KINEMATICS IN HEAD-FIXED MICE"

### headplate_1.png

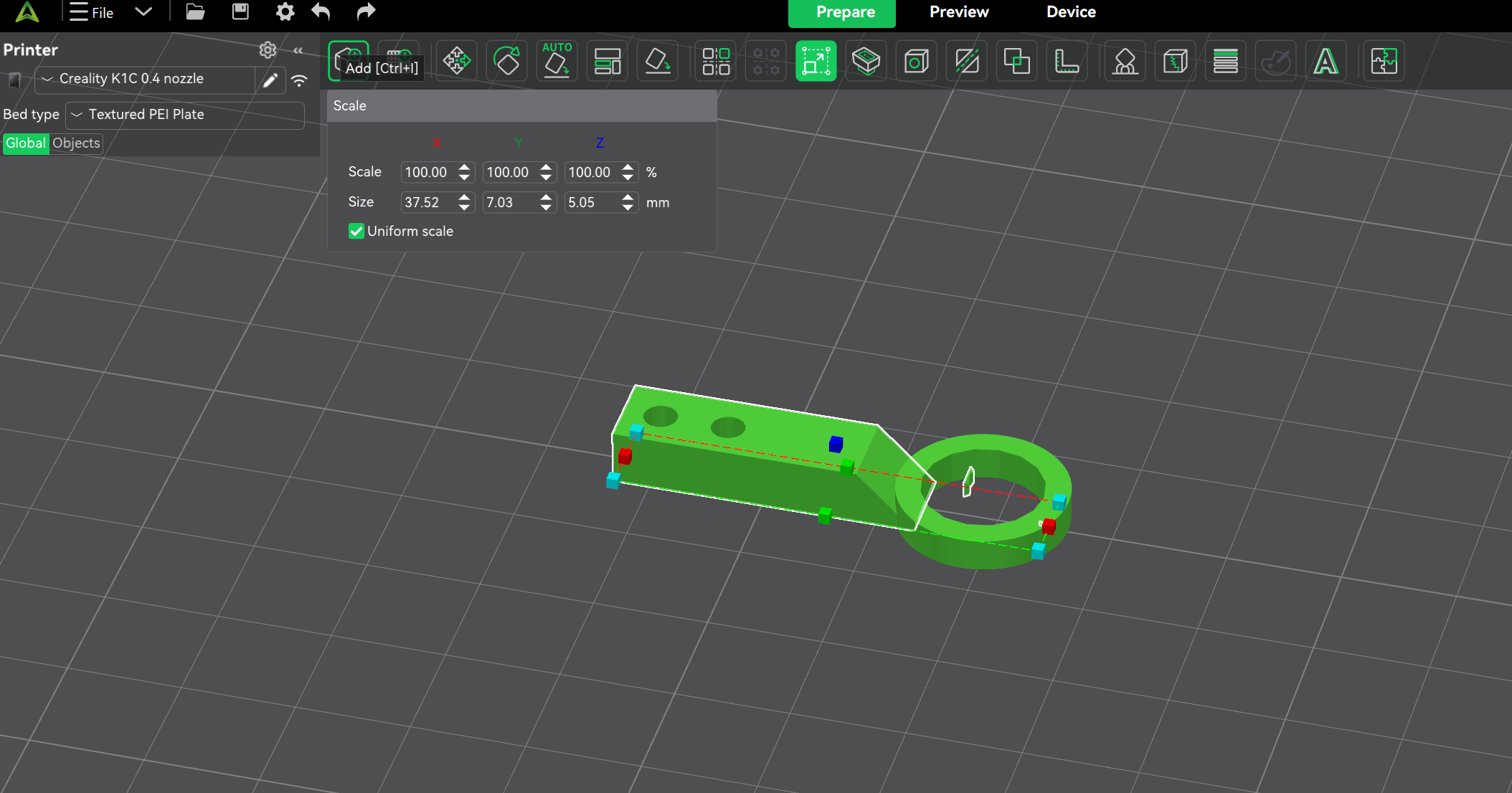

### headplate_2.png

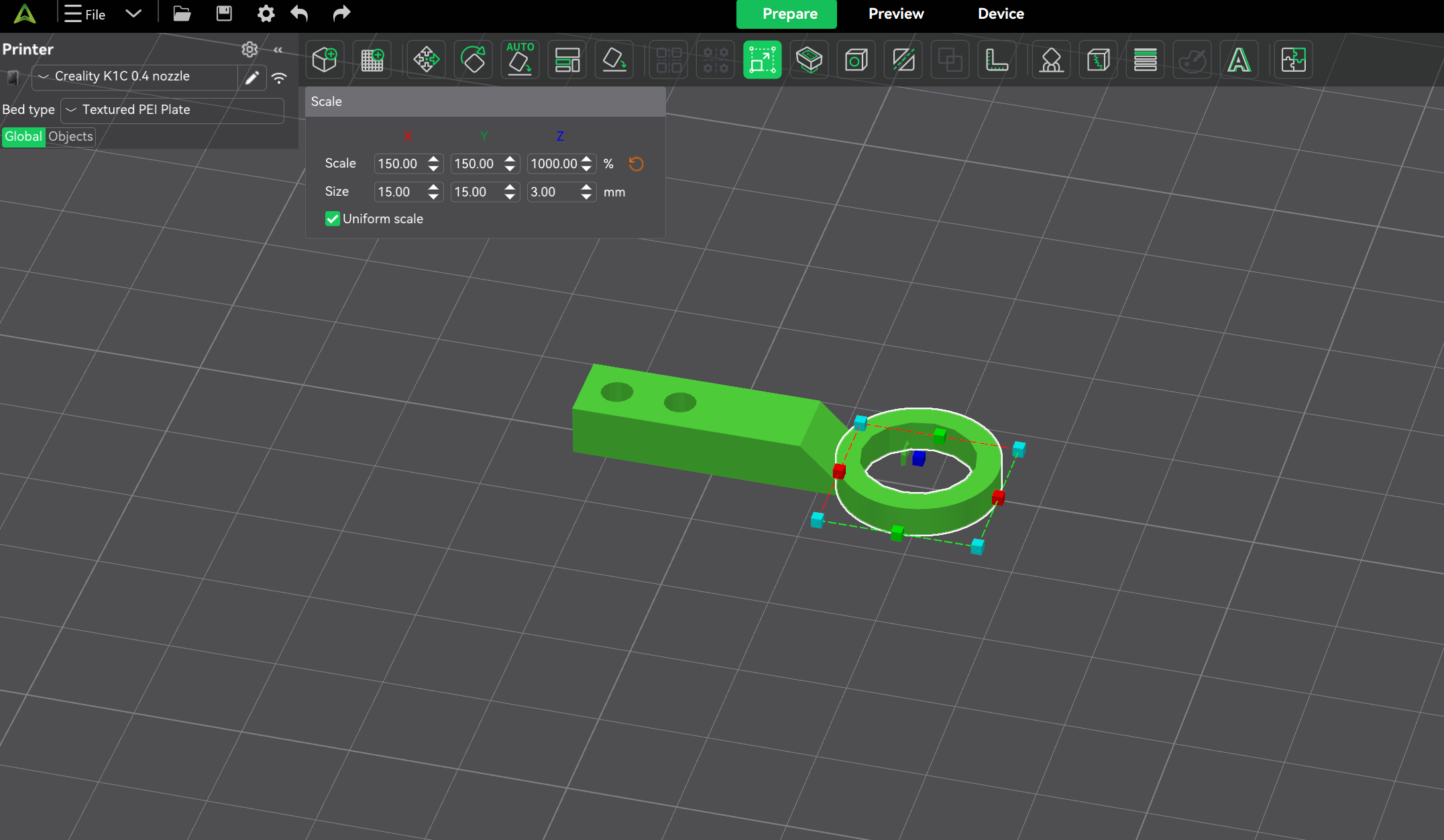

### headplate_3.png

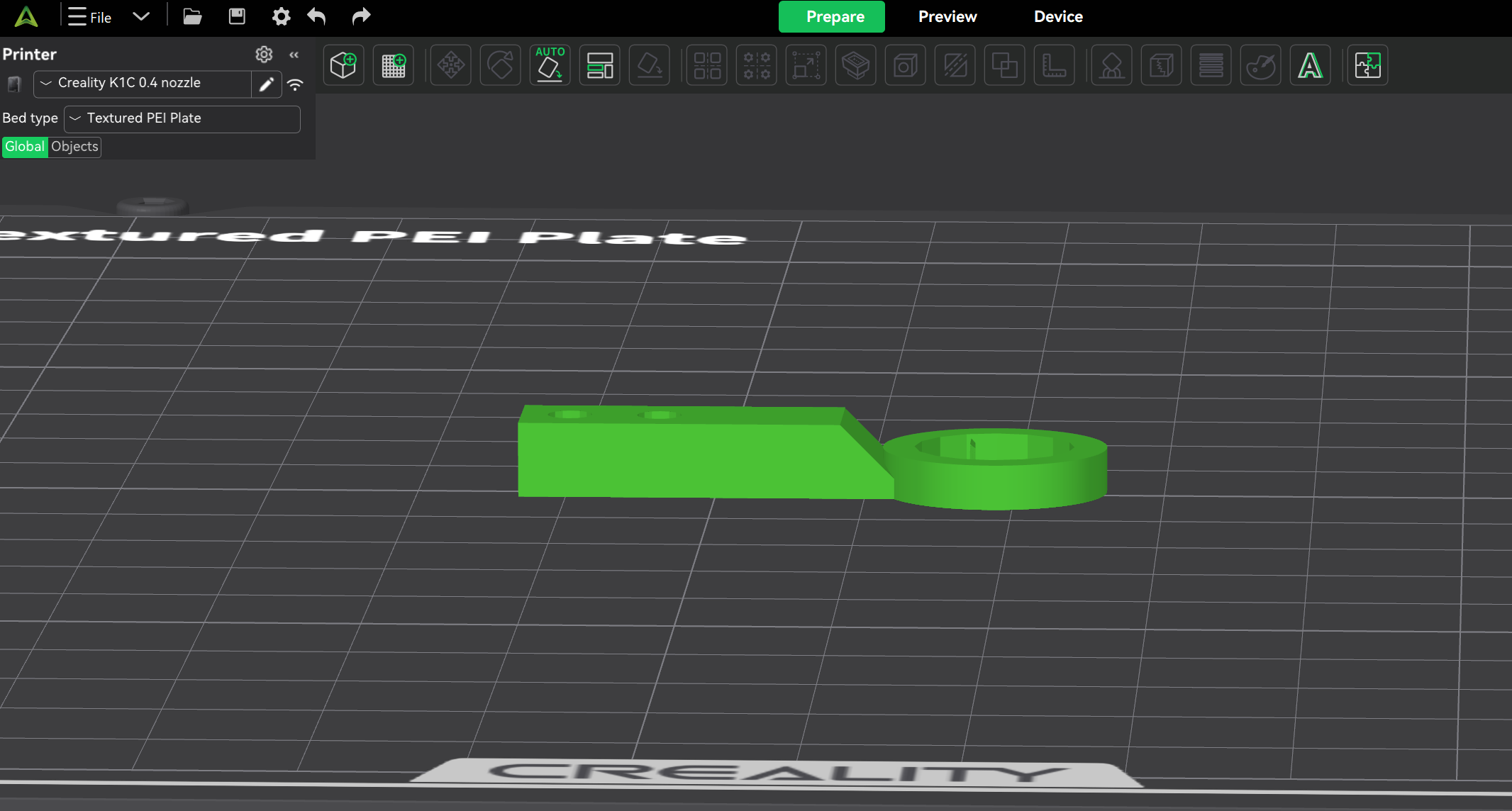

### headplate_4.png

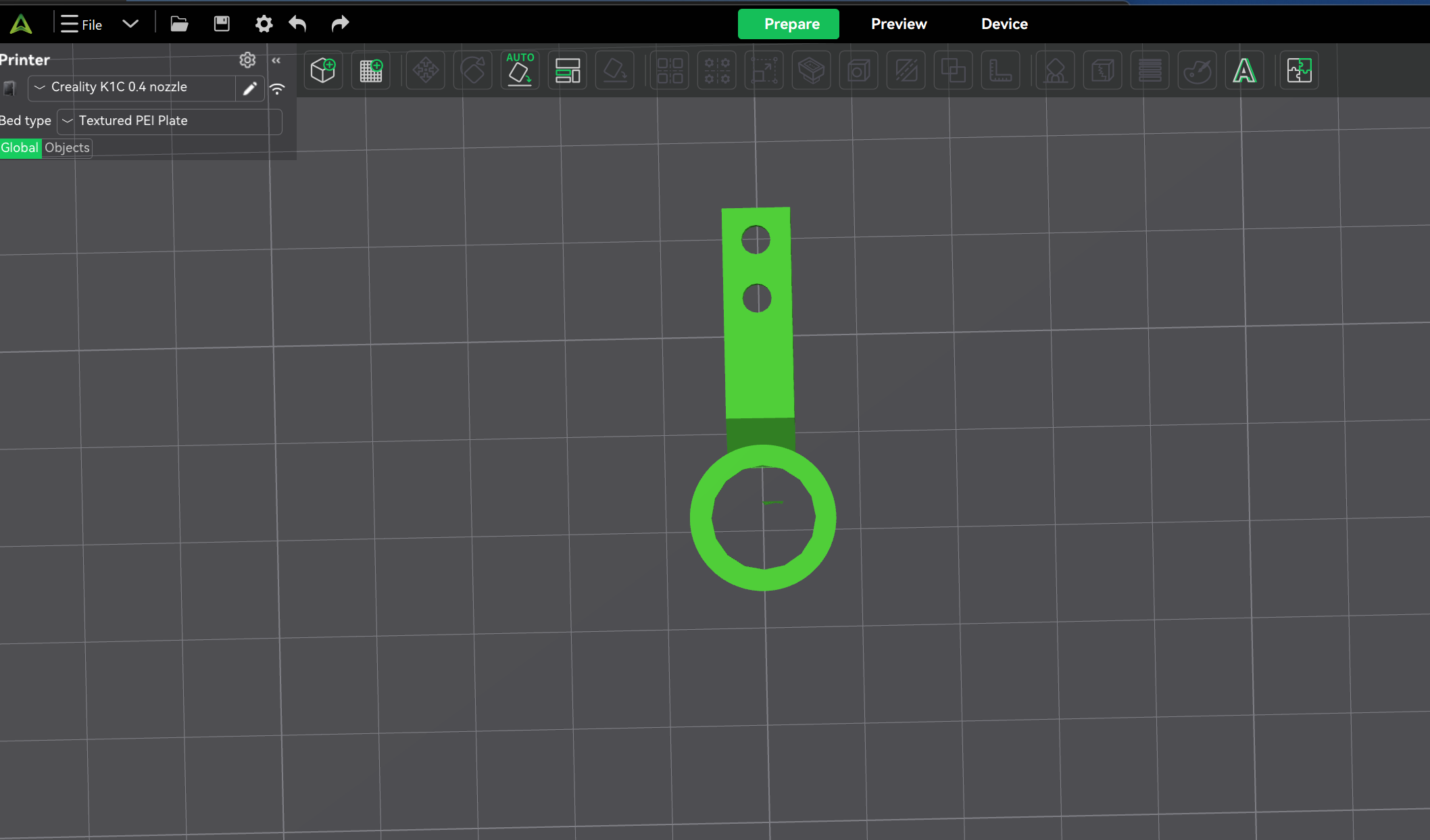

### headplate_5.png

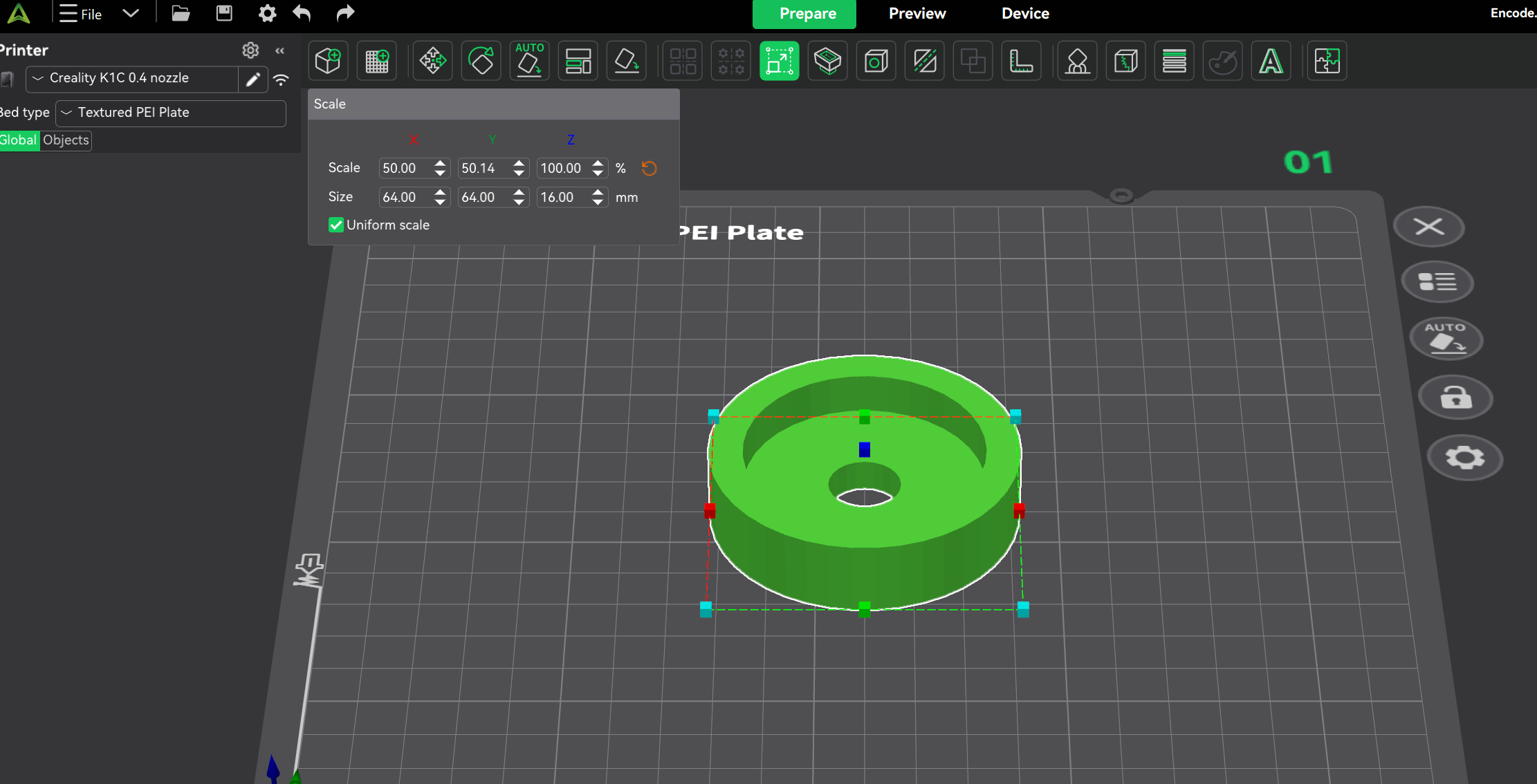

### headplate_6.png

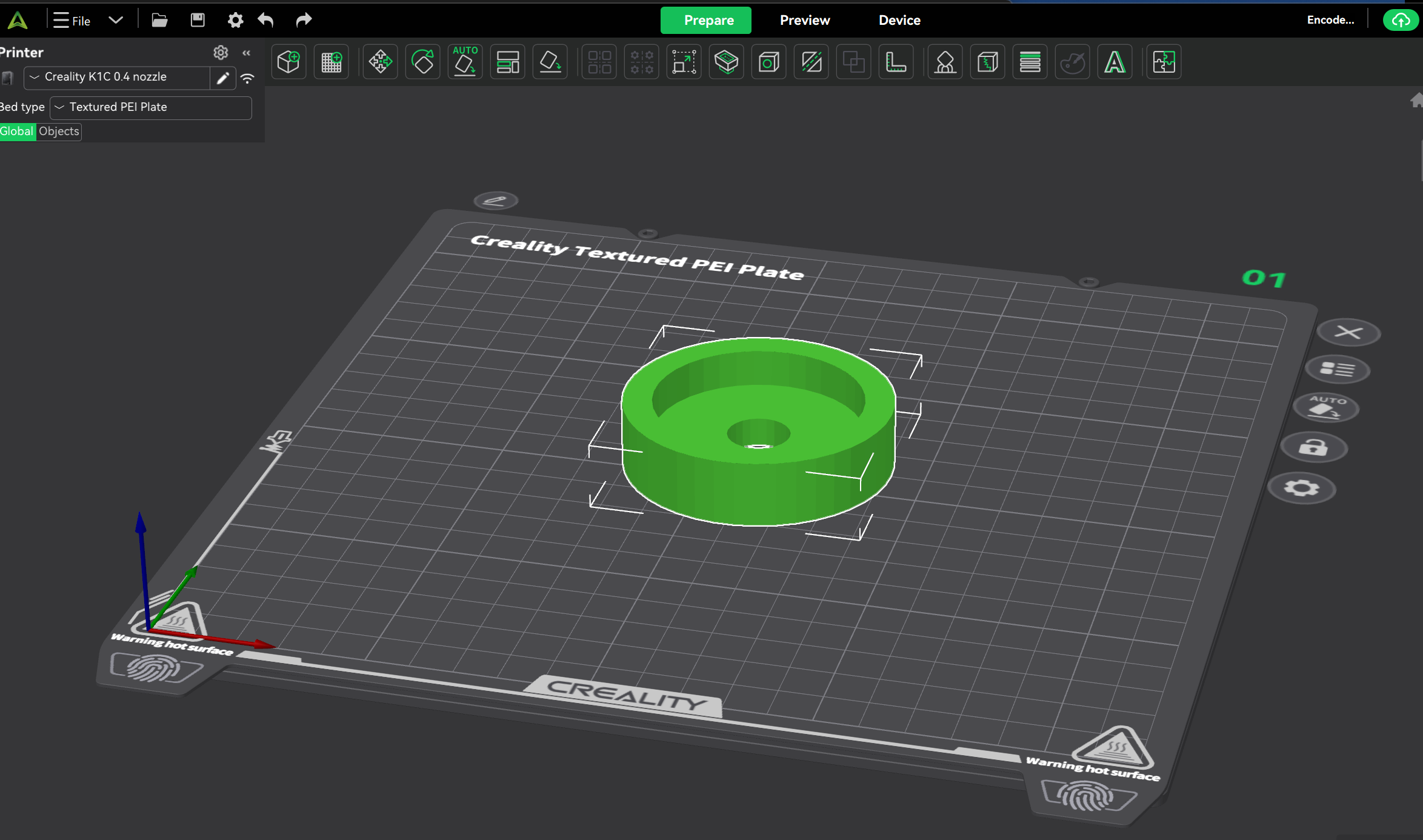
